# Pervasive context-dependent effects in the genetic architecture of complex and quantitative traits revealed by a powerful multiparent mapping population in yeast

**DOI:** 10.1101/2025.10.27.683165

**Authors:** Gareth A. Cromie, Russell S. Lo, Lauren Ames, Trey S. Morgan, Katherine Owens, Anne E. Clark, Martin S. Timour, Julee Ashmead, Michelle Tang, J. Nathan Kutz, Joshua M. Akey, Aimée M. Dudley

## Abstract

The genetic dissection of complex traits remains a major challenge in basic and biomedical research, but is essential for understanding the molecular pathways that shape phenotypic variation and for developing predictive models of trait and disease susceptibility. Here, we leverage a novel multiparent mapping population of budding yeast, CYClones, comprising 9,344 haploid strains derived from eight genetically diverse founders, to identify quantitative trait loci (QTL) and systematically investigate the genetic architecture of growth rates across ten environmental conditions. In total, we identified 349 QTL (ranging from 19 to 49 QTL per growth condition) that explained between 61% and 100% of narrow sense heritability across traits. The high power and resolution of CYClones revealed that growth traits exhibited distinct, condition-specific genetic architectures with extensive allelic heterogeneity, where a QTL was the result of multiple tightly linked causal variants. We also observed pleiotropy among QTL with complex, trait-dependent allele effects that are also consistent with allelic heterogeneity. Genetic complexity varied widely, with some traits showing nearly Mendelian architectures, while others were highly polygenic. Introgressed loci played a prominent role in the landscape of growth rate QTL, including one that is localized to a 2.4 kb interval in the *PCA1* cadmium transporter and explains 72% of variation in cadmium resistance and another large-effect QTL that alters growth on galactose via complex non-additive interactions with the transcriptional regulator GAL3. In both cadmium and galactose conditions, we show that allelic variation at a small number of loci stratifies the population into regulatory or physiological subgroups, each with distinct genetic architectures, a phenomenon we term allele-dependent stratification. Collectively, our results provide novel insights into the genetics of growth rates in budding yeast, the architectural features of genetic complexity, and demonstrate that CYClones is a powerful platform for revealing the molecular basis of complex trait variation.

## Introduction

Delineating the genetic architecture of complex traits is a fundamentally important problem with broad significance in genetics, evolution, and personalized medicine [1]. A deeper understanding of the relationship between genotypes and phenotypes will help elucidate the biological mechanisms contributing to variation in complex traits and enable phenotypes to be accurately predicted from an individual’s genome sequence. Mendelian traits and diseases are already routinely predicted from DNA sequences, such as in newborn screening programs [2]. Extending this paradigm to the prediction of complex traits and disease susceptibility, through polygenic risk scoring, has the potential to revolutionize medicine. However, polygenic scores have a number of limitations that need to be overcome, including accuracy and portability across populations, before being broadly useful [3].

Although large biobank-scale cohorts, such as the UK Biobank [4] and All of Us Project [5], have facilitated powerful genome-wide association studies (GWAS) in humans, understanding the rules governing genotype-phenotype relationships in outbred populations, like humans, is challenging. Specifically, high levels of allelic diversity and linkage disequilibrium in humans make identifying causal alleles difficult, particularly rare variants with small effect sizes [6]. Additionally, context-dependent effects, including gene x gene and gene x environment interactions, have proven difficult to assess in humans. Thus, model organisms have been a useful complementary strategy to understand the genetic architecture of complex traits. Large mapping populations derived from crossing multiple divergent parents have emerged as a popular study design to investigate the fundamental characteristics of genotype-phenotype maps [7–10]. These multiparent study designs, modeled after the Mouse Collaborative Cross [11], are powerful because they capture a larger proportion of population diversity compared to simpler crosses of two parental strains, and thus have a genetic background that more closely recapitulates outbred populations, while avoiding complications from rare alleles and the confounding effects of population structure [12].

Budding yeast, with its rapid growth and compact, well-studied genome is a particularly attractive model organism for genetic mapping. Very large mapping populations are feasible [13–15], enabling causal alleles with even small phenotypic effects to be mapped with high resolution. Importantly, the ability to more easily identify causal alleles facilitates the generation and experimental testing of mechanistic insights. In particular, large yeast mapping populations are well suited to investigate how factors such as context-specific effects, allelic and locus heterogeneity, and pleiotropy contribute to the genetic architecture of complex and quantitative phenotypes.

To this end, we constructed the CYClones (<u>C</u>ollaborative <u>Y</u>east <u>C</u>ross clones) mapping population using a multiparent funnel cross, analogous to the Mouse Collaborative Cross. In total, the final mapping population of CYClones consists of 9,344 haploid strains with low-coverage whole-genome sequencing data obtained for each strain. More specific details about how the CYClone strains were constructed are provided in a companion paper (https://doi.org/10.1101/2025.10.15.682626). We measured quantitative growth rates for all CYClone strains in ten different media conditions that represent independent evolutionary, ecological, and clinically relevant environments and performed comprehensive genetic analyses to estimate heritability and map QTL for each growth phenotype. Our results show that CYClones provides high statistical power to identify loci with weak phenotypic effects (down to 0.4% of phenotypic variation) and to map loci to single-gene resolution. These results are consistent with simulations which predicted ≥95% power to identify variants with heritability ≥ 0.36% with a resolution of <7.9 kb kb (described in the companion paper: https://doi.org/10.1101/2025.10.15.682626). In total, we identified 349 QTL, and the high resolution afforded by CYClones facilitated the discovery of 37 putative causal alleles mapped to single-gene resolution. Beyond mapping, our analyses reveal that the genetic architecture of growth phenotypes varies widely, ranging from Mendelian-like traits dominated by single loci to highly polygenic traits shaped by many small-effect variants. We also uncover pervasive context dependence, where the effects of genetic variants shift depending on regulatory or physiological state. In addition, we identify pleiotropic loci that influence multiple traits and allelic heterogeneity, in which multiple tightly linked variants at a single locus contribute to phenotypic variation. Together, these findings provide a detailed and mechanistically grounded view of the architecture underlying complex traits in yeast.

## Materials and methods

### Media and Genetic Manipulation of Yeast

Unless noted, standard media and methods were used for the growth and manipulation of yeast. Unless specified, yeast strains were grown in YPD medium (1% yeast extract, 2% bacto peptone, 2% glucose), with 2% agar for solid plates. Nourseothricin (NAT) was used at a final concentration of 100ug/ml. Hygromycin (HYG) was used at a final concentration of 500ug/ml.

### Condition Specific Growth of the Mapping Population

Yeast growth assays were carried out on solid media on Nunc™ OmniTrays™. With the exception of the galactose condition, all media were YPD (2% glucose) agar supplemented with the appropriate stock solutions (Table S1: https://doi.org/10.5281/zenodo.17362654). Galactose medium was YPGal solid medium containing the respiratory inhibitor antimycin A (1% yeast extract, 2% bacto peptone, 2% galactose, 1µg/ml antimycinA). Plates contained 35 ml of agar media dispensed by an automated plate pourer (Kreo Technologies). Plates were labeled with a computer-readable barcode to facilitate automated plate identification and dried at room temperature for three days before use. Each media condition was assayed twice, with biological replicates being performed on different days and with independently generated source plates.

To generate the source plates for replica pinning, the 96-well glycerol stock plates were thawed and 2.5 µl of each well was transferred to a fresh 96-well source plate containing 150 µl YPD broth per well. A control plate (used to estimate the effect of well position on patch growth) was inoculated with strain YO2685 in all 96 wells. Inoculated source plates were covered with a breathable seal (Agygen®, BF-400-S) and incubated at 30°C for 48 hours, without shaking. Following incubation, breathable seals were removed, and cells were resuspended by placing the plates in an orbital shaker at 900 rpm for two minutes. A Biomek i7 (Beckman Coulter) robot fitted with a 96-prong pinner (V & P Scientific, FP6 pins) was then used to spot strains from each source plate to one agar destination plate per experimental condition, shaking the source plate for five seconds between transfers. Once dry, pinned agar destination plates were placed in a 30°C incubator for 72 hours. Plates were briefly removed from the incubator and photographed every 24 hours using a mounted Canon PowerShot SX10 IS digital camera under consistent lighting, camera-to-subject distance, and zoom. Images (ISO200, f4.5, 1/40 s exposure) were acquired as JPEG files. The area (in pixels) and average grey scale value of each replica-pinned patch was extracted with PyPlate, a custom python pipeline built with functions from the scikit-image library (https://github.com/lacyk3/pnri-projects/tree/Image-Analysis-Demos/Funnel%20Cross%20Project).

Using PyPlate, each plate image was cropped into 96 regions of interest (ROIs) centered at the pin locations, with each region containing a single patch. Patch area was measured by segmenting each ROI using a combination of intensity thresholding and circle detection. Intensity thresholding was performed by binarizing the image with the Otsu threshold [16] and applying the morphological operations of *closing* and *filling* to segment the bright patch from the dark background. If the threshold returned by Otsu’s method was sufficiently different from the average background agar intensity, the results of Otsu thresholding were considered meaningful. Circle detection was performed by binarizing the image with an adjusted mean threshold and applying the Circular Hough transform [17] with a minimum radius equal to the initial pin radius. This exploits the geometry of the pinned cells to capture any regions of patches where growth is too faint to be detected with intensity thresholding. If the results of the intensity threshold were considered significant, these were combined with the detected circle to form the patch mask. However, if the Otsu threshold was similar to the background agar intensity, then the intensity thresholding was discarded and the detected circle alone was used as the patch mask. An additional check was performed to rule out unreliable results which may have arisen from a blank or contaminated ROI. Specifically, if the mean intensity of the masked patch area was too low or the likelihood of the detected circle was too low, then the patch was determined to be empty and assigned a value of NA. The final patch area is simply the total number of pixels in the patch mask.

### Phenotype Data Processing and Normalization

For 9,329 strains (99.8%) we obtained growth data from both replicates. Data was obtained from a single replicate for 15 strains. The 7 strains for which phenotyping failed in both replicates were removed from further consideration. The growth data were then normalized.

First, area measurements <=2000 pixels (approximate pin deposition area) were replaced with NA values. Along with patches that the image processing pipeline failed to identify, this left 0.3% of measurements as missing data (NA calls). Plate-level normalization was then carried out assuming a multiplicative plate effect, estimated using the 8 founder strain control patches on each plate. After this, multiplicative batch level normalization was carried out, where batches are groups of experimental plates processed in the same week. This normalization utilized all strains, equalizing the median area value for each timepoint + condition across batches. Finally, multiplicative normalization for growth on the four plate edges was carried out assuming a single timepoint-specific multiplier for each edge, adjusted by a condition-specific multiplier (across all edges) for each condition at each timepoint. Final phenotypes were calculated as the mean of all normalized measurements for each strain on each condition at each timepoint (Table S2: https://doi.org/10.5281/zenodo.17362654).

After these steps, the final dataset included 9,344 haploid (and euploid) strains haplotyped at 276,774 SNP positions and with normalized growth measurements at three time points (24, 48 and 72 hours) across 10 media conditions.

### Broad and Narrow Sense Heritability

Broad sense heritability was calculated as *H^2^* = 1−(σ^2^_e_/σ^2^_s_), where σ^2^_s_ is the variance of the population of segregant means, and σ^2^_e_ is the variance of the sampling distribution of these means. We calculated σ^2^_e_ based on the variance of duplicate measures of the same strain, using a random effects ANOVA implemented using the ‘lmer’ function in the R package “lme4” [18].

Narrow sense heritability was calculated using the R package “rrBLUP” [19]. First, a covariance matrix, K, was calculated using the ‘A.mat’ function applied to a matrix of allele values for all strains at the set of scaffolding markers. To produce the allele matrix, haplotypes were converted back into binary allele calls, with major alleles encoded with the value “1”, and minor alleles with the value “0”. Next, the ‘mixed.solve’ function was used with the K matrix and the vector of observed phenotypes for each condition at each timepoint to calculate additive (V_a_) and residual (V_e_) variance components. The narrow sense heritability for each condition at each timepoint was then estimated as V_a_/(V_a_+V_e_). Standard errors were then calculated as the standard deviation of 100 bootstrap replicates of each time + condition.

### QTL Mapping

For single-marker linkage mapping, at each tested position, the LOD score was calculated as LOD=n/2 x log_10_(RSS_0_/RSS_1_), where RSS_0_ is the null residual sum of squares assuming no QTL (single population mean) and RSS_1_ is the residual sum of squares assuming a QTL (fitting a separate mean for the 8 subpopulations defined by the haplotype at the current marker).

For pairwise-marker linkage mapping of genetic interactions, at each tested pair of positions (*s*,*t*) the LOD*_i_* score, measuring the improvement in the fit of a model allowing interaction (L*_f_*) over an additive model (L*_a_*), was calculated as L*_f_*(*s*,*t*) – L*_a_*(*s*,*t*), where L*_f_*(*s*,*t*) and L*_a_*(*s*,*t*) are the log_10_ likelihoods for the interaction and additive models, respectively. Likelihoods were calculated from the residuals of each model, assuming normality.

We used permutations to determine a LOD score threshold that corrected for both the number of markers tested in each genome-wide scan and the total number of phenotypes analyzed. Specifically, we carried out 1000 permutations of which 10-phenotype vector (from the 72h dataset) was associated with each haplotype vector. We then carried out single marker linkage analysis as above and identified the highest LOD score across all phenotypes for that permutation. The 95^th^ percentile of the 1000 permutation maximum LOD values was used to define a genome-wide, phenotype-corrected 5% FWER LOD score of 8.3.

Because estimating family-wise error rates for the pairwise marker analyses was computationally intensive, instead a 1% false discovery rate (FDR) was calculated for each of the pairwise non-additive datasets using a null distribution for each condition at each timepoint. The null distribution was generated by calculating the interaction LOD, as above, using a single permutation of the residuals from the additive model as the phenotype for each pair of markers.

### Generation of Additive Genetic Models and Fine Mapping of QTL

For each condition and timepoint, additive genetic models were constructed by an iterative process using forward selection. In the first iteration, single-marker LOD scores were calculated for each of the 5479 scaffolding markers, and we retained the most significant marker per chromosome if it exceeded a LOD of 9, conservatively based on the 5% FWER LOD score threshold described above. Then, a linear model was fit using the phenotype values and the haplotypes at the set of identified markers. The residuals from this model were then used as the phenotype values and a second genome-wide linkage analysis was performed, identifying the most significant marker per chromosome, as above. The markers identified from this analysis were then added to the existing set of markers and used to fit a linear model to the phenotype values. We repeated this process until no markers above the LOD score threshold were identified or until an ANOVA, comparing the model from the current iteration to the preceding model, failed to show a significant improvement (p-value>0.001).

We next fine-mapped the position of each QTL and generated confidence intervals of the location of each QTL. For each marker in each final model, that marker was removed from the model and the residuals of the model were adjusted to reverse the phenotypic effects attributed to that marker. Then, linkage mapping using the adjusted residuals was carried on the same chromosome as the “dropped” marker but using all markers (from the full set of ∼270k), not just markers from the scaffolding set (∼5.5k). The strongest marker was then identified along with a 2-LOD confidence interval. Finally, genes were identified as falling within the confidence interval of each QTL if the interval overlapped a region extending from 1kb upstream to 0.5kb downstream of a gene’s open reading frame.

QTL traces were highly similar across the three timepoints of our experiments (Table S3: https://doi.org/10.5281/zenodo.17362654). Based on this, the confidence intervals of the 24h and 48h QTL sets for each condition were used to further refine the gene-level resolution of the QTL measured at the 72h timepoint. For each 72h QTL, the list of genes falling within its 72h confidence interval was compared to the list of genes falling within any QTL confidence interval in the 24h and 48h datasets for the same condition. Only QTL with confidence intervals <100 kb were refined or used for refinement. The genes with the highest (or co-highest) count were retained. Because the set of genes lying within the set of QTL confidence intervals is always a small subset of all genes, this approach effectively examines overlap between confidence intervals for the same QTL at multiple timepoints without having to first estimate the true physical position of the (shared) QTL.

### Variance Explained by Additive and Interaction Models

The variance explained by additive models from each timepoint and condition was calculated by 10-fold cross-validation. In each fold, the training data was used to generate an additive model by forward selection, as described above (without fine mapping), and the variance explained by that model was calculated on the test data. The mean test-population variance explained across the 10-folds was then calculated.

To examine genetic architecture, we then took the models from each training fold and ordered the markers from the most to the least significant (type II ANOVA p-value) based on the training fit. Next, in each fold, we fit a new model using only the single strongest marker from the training data, measured the variance explained in the corresponding test fold, and took the mean variance explained across all folds. This number represents the variance explained by the strongest single marker model. We then repeated this using the strongest two markers from the training models. This process was continued until the number of markers reached the size of the smallest full model in any of the training folds.

We also took each of the full additive models from the 10 folds and introduced significant non-additive interactions between the loci, in a stepwise fashion. In each training fold, for each pair of QTL, a new model was constructed identical to the additive model except for allowing a non-additive interaction between the two current loci. Each of these models was compared to the base additive model by ANOVA, and ranked based on the resulting p-value. The additive model was then extended, one pairwise interaction at a time, using the rank-ordered pairwise interactions (strongest to weakest). After each model update, the new model was compared to the previous iteration by ANOVA and the process was stopped if the improvement was not significant (p-value<0.001). The average variance explained by the final models was calculated on the corresponding test folds as above.

### Determination of QTL Multimodality

For each 72h QTL identified in the fine-mapping additive models, the 8 haplotype effect size estimates were extracted. Estimates associated with haplotypes seen in <50 strains were replaced by NA values. K-means clustering (k=2) was applied to the effect size estimates for each QTL in order to binarize the haplotypes. Then, for each QTL, 100 random 90:10 training:test strain splits were generated. For each training set, three models were fit, the first using the full haplotype encoding at the current QTL (full), the second the binary recoding of the current QTL (binary), and the third lacking the current QTL (null). Each model incorporated (using the original haplotype encoding) all other QTL in the fine-mapping additive model. For each test set, the additional variance explained (relative to the null model) was then calculated for the full and binary models. The distributions of these values were used to calculate whether the full models explained significantly more variance than the binary models (one-sided t-test, p-value<0.05 after Holm correction for all QTL tested across all conditions). The means of the distributions were used to estimate the variance explained by the full and binary models for each marker.

### Identification of Pleiotropic Genes

To identify QTL mapped to single gene resolution affecting more traits than expected by chance, the number of the single genes mapped in each condition was counted and the identity of these genes was randomized (across all yeast genes) and the maximum number of conditions sharing a gene was calculated for each permutation of all conditions. A total of 10,000 permutations was carried out and the 95% quantile of this null distribution was identified as 2 conditions, so that a gene shared by 3 or more conditions is significant at a p-value<0.05

### GAL3 Variant Construction and Testing

The *GAL3* region of founder strains YO2685 and YO2563 was amplified from genomic DNA and cloned into the pRS41N plasmid,^19^ via Gibson assembly. Single nucleotide changes in *GAL3* were then induced by Quikchange PCR. All resulting plasmids (Table S4: https://doi.org/10.5281/zenodo.17362654) were fully sequenced by Oxford Nanopore Sequencing. Plasmids containing *GAL3* variants were transformed into either YO2685 or YO2563 and growth on YPGal medium was quantified, using PyPlate, from images of yeast strains replica pinned onto solid agar plates.

### GAL Statistical and Subpopulation Analyses

The interaction versus additive model values for each scaffolding marker in combination with *GAL3* were taken from the full pairwise interaction versus additive model matrix, using the row corresponding to the peak of the *GAL3* QTL, marker chr04_465458.

Because of the strength of the *GAL3* QTL, for interaction and subpopulation analyses, strains with crossovers between the markers flanking GAL3 ("chr04_463411” and “chr04_465458”) were removed before further analysis, as their *GAL3* QTL haplotype is ambiguous. Similarly, haplotype 6 identifies the introgression at the 3-gene *GAL1/7/10* locus, so strains with haplotype 6 at only one of the markers flanking this locus (“chr02_271559” and “chr02_281559”) were also removed. The *GAL1/7/10*, *GAL2*, *GAL3*, *GAL4*, *GAL80* and *PGM1* loci were specified using the markers “chr02_271559”, “chr12_292952”, “chr04_463411”, “chr16_80216”, “chr13_171614”, and “chr11_203623”, respectively.

To test the significance of the four-way non-additive interaction linear model between *GAL3* and introgressed vs reference alleles of *GAL1/7/10*, *GAL2* and *PGM1*, *GAL3* haplotypes were first discretized into two categories (*GAL3*^+^ vs *GAL3*^-^). An interaction model involving all four genes was then compared, by ANOVA, to a linear model with an additive effect of *GAL3* and interactions between the other three genes.

For estimating the variance explained by markers in the *GAL3*^+^ vs *GAL3*^-^ subpopulations (both lacking the *GAL1/7/10* introgression), 100 random training/test strain splits (90:10) were carried out for each marker in each subpopulation. For each marker, an additive model was fit on the training subset of each split. Then the model was applied to the matching test dataset and the residual phenotypic variance was calculated. The mean variance explained and the associated standard error was then calculated across the splits.

For each GAL subpopulation, the effectiveness of global and subpopulation-specific additive models was assessed as follows. For each of the subpopulations (*GAL3^+^*, *GAL3^-^*, *GAL1/7/10* introgression), defined as described above, 100 random training/test strain splits (90:10) were carried out. For each subpopulation and split, an additive model was fit to the training set as described above along with a null (mean only) model. The models were then applied to the matching test datasets and the residuals were calculated. For each of the 100 splits, a global model was also fit after first combining the training strains from each of the three subpopulations. This model was then applied to each test subpopulation individually, and the residuals were calculated.

### CAD Statistical and Subpopulation Analyses

For the major CAD peak on chromosome II, the QTL is close to the right telomere which has an introgression in founder 2. Because of sequence divergence between the introgression and the S288c reference genome, read pairs from strains harboring the introgression (haplotype 2) frequently fail to align to the reference genome, causing undercounting of haplotype 2 at variant sites across this region. To more accurately infer the haplotype this region, read-pairs from each strain that did not align to the S288c reference genome were aligned, using the same parameters used in the original alignment to the reference genome, to the sequence of the right end of chromosome II from founder 2 (chromosome sequence assembly by Plasmidsaurus), corresponding to positions 750,001+ on the reference sequence (732,712+ in the founder 2 chromosome assembly: File S1: https://doi.org/10.5281/zenodo.17362654). The positions of variant sites from the original sequencing were mapped onto the introgression reference and counts of minor and major alleles were updated using the newly aligned reads as described for the S288c reference sequence, above. Haplotyping of chromosome II was then repeated using the previously described haplotype inference methods. The old and new chromosome II haplotypes were then merged for each strain, replacing the old haplotypes from the right end of chromosome II to the first variant position greater than 750,001 where the original and new datasets call the same parental haplotype. Relative to the original haplotype, some markers are lost from the extreme right telomeric region of chromosome II, as this region is not present in the introgressed sequence. Strains were then split into *PCA1* subpopulations based on their haplotype at scaffolding marker “chr02_794058”.

For both the GAL and CAD subpopulation analyses, QTL mapping was carried out as described above. For each mapped (sub)population, a 1% significance threshold was calculated from a null distribution generated from 1000 permutations of the data. In each permutation, the haplotype-phenotype assignments were randomized and the maximum LOD score was calculated for the (single) phenotype of interest. Additive models were fit as above.

## Results

### Overview of the CYClones mapping population

To construct the CYClones mapping population, we selected eight haploid “founder” strains isolated from distinct geographic and environmental sources (Fig 1A) and crossed them in a funnel design (Fig 1B). The eight parental strains were carefully selected to capture a significant proportion of segregating variation in the global budding yeast population (Fig 1A and 1B). Specifically, the eight parental strains are genetically diverse, with ∼270k high-quality single nucleotide variants (SNVs) (Table S5: https://doi.org/10.5281/zenodo.17362654), a density of ∼1 SNVs per 44 bases, that include 56% of common variants (minor allele frequencies greater than 0.05) and 32% of low-frequency and rare variants (minor allele frequencies between 0.05 and 0.005) in the global *S. cerevisiae* population [20]. Construction of the CYClones library employed three rounds of crossing (Fig 1B) to produce 11,484 haploid strains that are recombinant mosaics of the eight founder haplotypes. Unlike mice or other diploid organisms, these haploid yeast strains did not require further inbreeding to achieve homozygosity. Low coverage whole genome sequence analysis with imputation was used to genotype the ∼270k founder SNVs in each segregant (Fig 1C and Table S6: https://doi.org/10.5281/zenodo.17362654) and to identify strains with aneuploidies. This resulted in a final mapping population of 9,344 fully genotyped euploid, haploid strains. A detailed description of the construction of the CYClones population is provided in a companion publication (https://doi.org/10.1101/2025.10.15.682626). Simulations described in that paper indicate that CYClones is expected to have ≥95% power to identify variants with even very weak phenotypic effects (heritability ≥ 0.36%) at a resolution of a few kb or less, which is often less than the length of a single gene.

**Fig 1.**
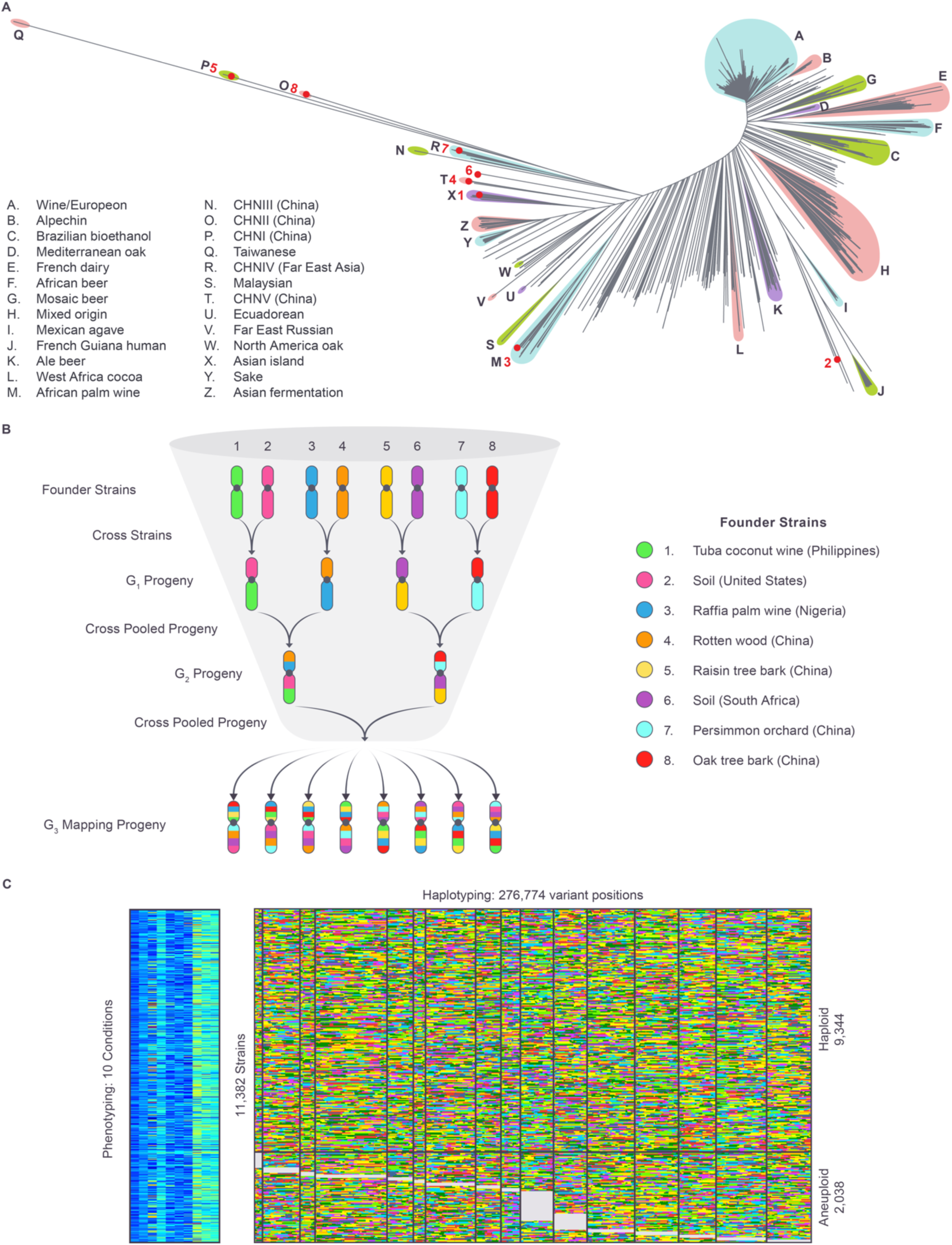
Mapping population and dataset. (**A**) Phylogenetic tree of the global *S. cerevisiae* population with subpopulations from Peter *et al*. [8] and the eight funnel cross founder strains (numbered 1:8 in red) indicated. (**B**) The eight founder strains were used to generate a mapping population via a funnel cross design. Each founder strain was mated pairwise to produce heterozygous diploids, which were each sporulated to produce pools of 576 MAT**a** and 576 MATα haploid progeny. Two rounds of reciprocal crossing using pooled strains were then carried out to produce the final mapping population of 11,484 strains. (**C**) Each of the strains in the final mapping population was individually genotyped and karyotyped by low coverage WGS. Growth of each strain on agar plates was measured on ten different media, in duplicate, and at 3 time points. After quality control filtering, this resulted in a final population of 9,344 euploid and 2,038 aneuploid strains each individually haplotyped, with growth phenotypes measured individually on 10 media conditions. Haplotype colors as in **B**.

### High precision phenotyping for ten independent growth conditions

For each of the 9,344 euploid strains, we measured growth rates on ten different media conditions that perturb a diverse set of biological processes representing evolutionary, ecological, and clinically relevant environments (Table S2: https://doi.org/10.5281/zenodo.17362654). The conditions tested were: rich medium containing the carbon source galactose (GAL), rich medium containing the carbon source glucose (YPD), YPD supplemented with the antifungal drugs caspofungin (CF) or fluconazole (FLZ), and YPD supplemented with the chemical stressors cadmium chloride (CAD), caffeine (CAF), calcium chloride (CAL), hydroxyurea (HU), rapamycin (RAP), or sorbitol (SOR). These conditions sample a broad range of physiological, stress, and ecological perturbations that natural populations of budding yeast might encounter, allowing us to explore the general principles of the genetic architecture of quantitative trait variation. For phenotyping, strains were robotically pinned in duplicate from liquid culture onto solid-media plates. Growth at three timepoints (24, 48 and 72 hours) was then assessed from colony area by automated image analysis (Methods). Reproducibility was very high, with a median *r^2^*value of 0.93 between the independent measurements for each condition at 72h. As was seen in a previous study measuring the effect on growth of all viable yeast gene deletions under many of the same conditions [21], growth values on the chosen conditions were largely uncorrelated, with a median *r^2^* value of 0.04 (Table S7: https://doi.org/10.5281/zenodo.17362654), suggesting that distinct biological processes are required for growth under these conditions.

### Highly diverse genetic architectures characterize the ten traits

Phenotypic variation was overwhelmingly driven by genetics for all ten conditions, with the estimated proportion of the phenotypic variance explained by genetic effects (broad-sense heritability, H^2^) ranging from 95-99% at 72h (Fig 2A) and >90% at all timepoints. As such, this dataset should be highly powered to dissect genotype-phenotype associations. We next estimated the proportion of the phenotypic variance explained by purely additive genetic effects (narrow sense heritability, h^2^). The variability of this metric, ranging from 38% (rich medium, YPD) to 76% (cadmium chloride, CAD) at 72h (Fig 2A), suggested that diverse genetic architectures underlie the ten traits being studied. To characterize these genetic architectures, we performed haplotype-based linkage mapping for each condition at each timepoint using a “scaffolding” subset of 5,479 markers (out of the total set of ∼270k markers) spaced approximately 2 kb apart across the genome (Table S3: https://doi.org/10.5281/zenodo.17362654). Simulations demonstrated that this dense set of scaffolding markers shows very little loss of sensitivity for QTL detection relative to the full set of ∼270k markers (described in the companion paper: https://doi.org/10.1101/2025.10.15.682626). As the results were highly similar across timepoints, we focused on the 72h results. This analysis confirmed the power of our dataset, which was able to detect QTL effects explaining as little as 0.4% of phenotypic variation (5% family-wise significance: LOD=8.3), a result consistent with simulations which predicted ≥95% power to identify variants with heritability ≥ 0.36% (described in the companion paper: https://doi.org/10.1101/2025.10.15.682626). The ten traits examined were all highly complex, with multiple significant QTL peaks identified in each condition. As expected, substantial variation was observed in the genetic architectures underlying the different traits (Fig S1 and Table S3: https://doi.org/10.5281/zenodo.17362654). Not only did the positions of the QTL peaks vary between conditions, consistent with the trait-to-trait differences in which genes are sensitive to genetic variation, but the growth phenotypes exhibited markedly distinct degrees of complexity. Some phenotypes, such as growth in GAL (Fig 2A and Fig S1) displayed an almost Mendelian architecture, dominated by a single QTL (max LOD = 1101; 42% of total variance), whereas other conditions, such as FLZ, SORB and YPD were highly polygenic with no single locus explaining a high fraction of phenotypic variation (Figs 2A, 2B and Fig S1).

**Fig 2.**
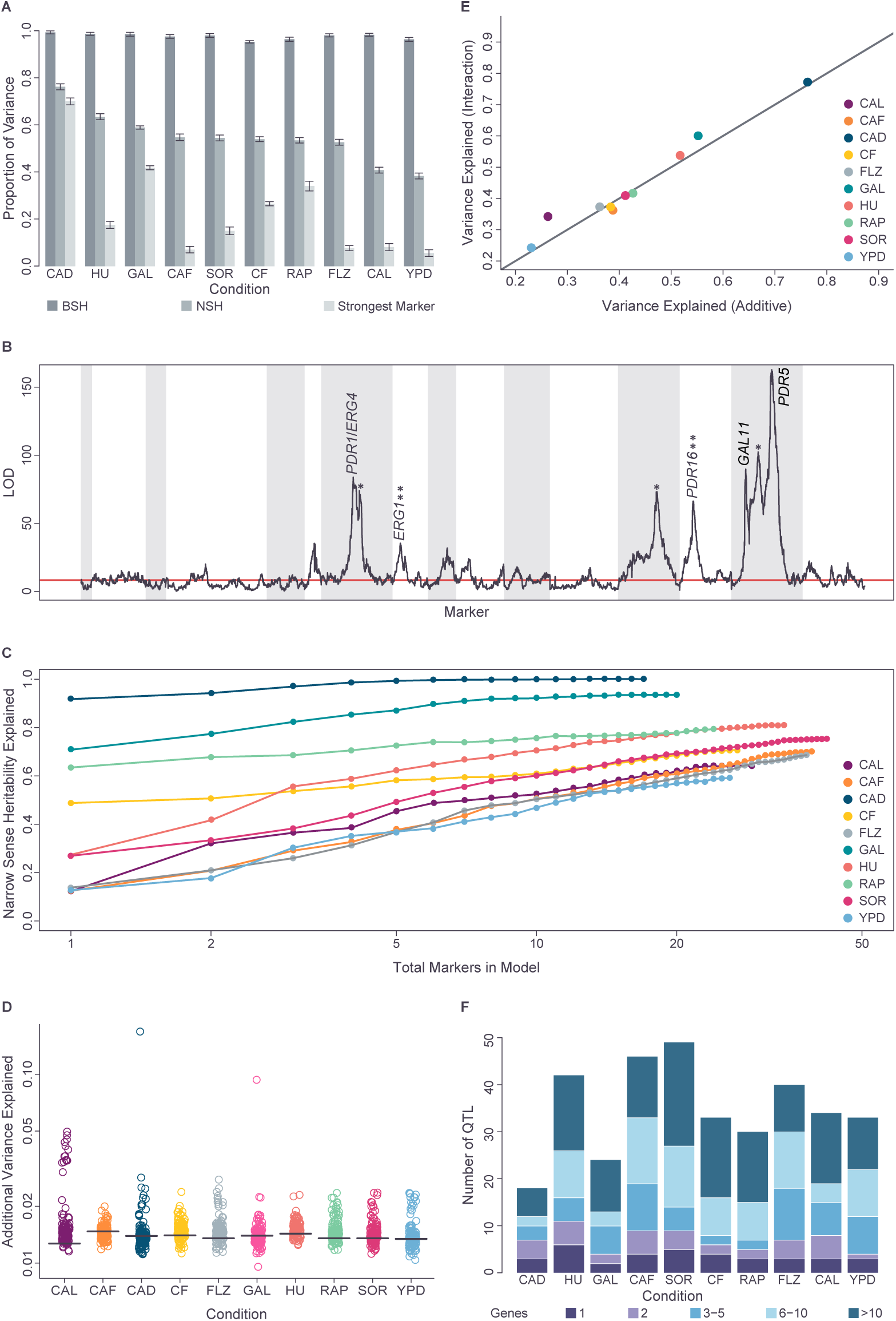
Genetic architecture of ten traits. (**A**) Broad (BSH) and narrow (NSH) heritability, and variance explained by single strongest marker, for all conditions at 72h. (**B**) Single scaffolding marker LODs for fluconazole, chromosomes alternating grey and white. Known resistance genes mapped to 1, 2 or 3-5 gene (**) resolution are indicated. Single asterisks indicate non-FLZ-specific peaks. (**C**) Proportion narrow sense heritability explained by additive models as QTL are added in order of strength. (**D**) Maximum additional proportion variance beyond additivity explained by inter-chromosomal pairwise interactions with 1% FDR thresholds (black horizontal lines). (**E**) Effect on total proportion variance explained of allowing non-additive interactions within additive models. (**F**) Additive model QTL number and resolution (number of genes in 2-LOD interval).

### Genetic models built using CYClones can explain most of the additive phenotypic variation across all phenotypes

To explore these architectures further, we developed an additive genetic model for each condition by forward selection using the scaffolding markers (Methods). For all traits, the additive models explained most of the narrow sense heritability (60-100% by 10-fold cross validation) highlighting the power of the mapping population. Our results are consistent with Fisher’s infinitesimal model [22], with QTL number increasing as effect size decreases (Fig 2C), similar to the results of a recent large-scale budding yeast Barcoded bulk QTL study [23]. However, for several of the conditions (e.g. YPF, FLZ) there is a substantial gap between narrow and broad sense heritability (Fig 2A), indicating that, for these traits, genetically-determined phenotypic variation cannot be fully explained by additive models and suggesting a role for non-additive genetic interactions.

Comparison of additive versus interaction models for all pairs of scaffolding markers in all conditions confirmed that our dataset is also highly powered to detect pairwise (non-additive) interactions, with a (1% FDR) threshold of ∼1.4% of variance explained (Fig 2D). In our dataset, weak, but significant, pairwise non-additive interactions were common, but strong non-additive interactions were rare (Fig 2D). Extending our additive genetic models to allow non-additive interactions between the markers in those models, showed little improvement in performance, as assessed by 10-fold cross-validation (Fig 2E). This pattern is consistent with results from both human GWAS and model organism studies, where additive models capture most heritable variance, and including pairwise interactions only modestly improves prediction [24–26].

### CYClones enables large numbers of QTL to be mapped at high resolution across diverse genetic architectures

For each trait, we were able to identify large numbers of statistically significant QTL from the additive models, from 18 in CAD to 49 in SOR, totaling 349 QTL across all conditions at 72h. These QTL were added to the model iteratively using a LOD cutoff of 9 to account for multiple marker and condition testing (Materials and Methods). These QTL were then mapped at high resolution using the full set of ∼270k high-quality variant sites, with 66 (19%) mapped to intervals of less than 5 kb and 173 (50%) mapped to intervals of less than 20 kb. Using the mapping data from all three timepoints to further resolve the position of each QTL (Methods), enabled 37 putative causal alleles to be mapped to single gene resolution in the 72h dataset. An additional 32 and 25 QTL were mapped to a resolution of 2 and 3 genes, respectively and, in total, 132 QTL (38%) were mapped to a resolution of 5 genes or less (Fig 2F and Table S8: https://doi.org/10.5281/zenodo.17362654). For conditions with well-characterized genetics, such as fluconazole-resistance (Fig 2C) and growth on galactose (below), mapped loci included known target genes resolved to the one- or two-gene level (Table S8: https://doi.org/10.5281/zenodo.17362654). The ability to accurately identify small numbers of genes, including single genes, underlying many QTL demonstrates that CYClones is a powerful tool for producing candidate gene lists highly enriched for causative genes, enabling the biological mechanisms underlying phenotypic variation to be elucidated.

### Heterogeneous allele effects characterize pleiotropic loci

We next leveraged the fact that the ten traits chosen for this study showed very little correlation (Table S7: https://doi.org/10.5281/zenodo.17362654) and displayed very distinct QTL landscapes (Fig S1 and Tables S2, S3, and S8: https://doi.org/10.5281/zenodo.17362654), as expected for traits that require distinct biological processes. As such, a causative gene impacting multiple conditions in our study would be consistent with the most conservative definition of pleiotropy, where a single gene impacts multiple distinct processes. We therefore tested whether our ability to map multiple QTL to single gene resolution would allow us to identify cases in which genes exhibited pleiotropic effects. To identify statistically significant instances of pleiotropy, we performed permutation analysis and determined that a gene mapped at single resolution in more than two conditions would be unlikely to occur by chance (p-value<0.01). This analysis identified three such genes: *FLO11* (*MUC1*), *HPF1,* and *WHI2*. *FLO11* encodes a flocculin involved in cell adhesion and biofilm formation was mapped to single gene resolution in five conditions and at two gene resolution in a further five conditions (ten conditions, total). *HPF1* encodes a cell wall mannoprotein and was mapped to single gene resolution in seven conditions. Finally, *WHI2* encodes a negative regulator of TORC1 and was mapped to single gene resolution in five conditions and at three gene resolution in another two conditions (seven conditions, total).

Pleiotropy can occur when alleles of a single gene have a consistent effect on multiple traits each having otherwise substantially different underlying genetics (i.e. uncorrelated traits). It can also occur when allele effects at a single locus differ between traits, for example, because multiple variants are present at the locus and these variants each affect different traits. Using our dataset, in the first case, we would expect to see correlated haplotype effects for a pleiotropic locus across different growth conditions. In the second case, we would expect to see uncorrelated haplotype effects. In fact, both patterns are seen across the shared loci identified in our study (Table S9: https://doi.org/10.5281/zenodo.17362654). For *HPF1*, a single haplotype effect pattern was seen across all conditions (Figs 3A, 3B and Table S9: https://doi.org/10.5281/zenodo.17362654) with the first principal component explaining 85% of the total variance of the standardized haplotype effects. In contrast, at *FLO11* and *WHI2*, some conditions were closely correlated with each other, while others showed different patterns of haplotype effects. Most dramatically, at *WHI2*, two distinct and opposite haplotype effect patterns were observed (Figs 3C, 3D and Table S9: https://doi.org/10.5281/zenodo.17362654). In CAL, HU, SOR and YPD, haplotypes 1 and 6 promoted growth relative to the other haplotypes, while in CAD, GAL and RAP these haplotypes inhibited growth relative to the others (Fig 3C and Table S9: https://doi.org/10.5281/zenodo.17362654). This pattern is consistent with variants at this locus having the same effect on gene activity in all conditions, but in some conditions the gene activity has a positive effect on growth, while in other conditions it has a negative effect.

**Fig 3.**
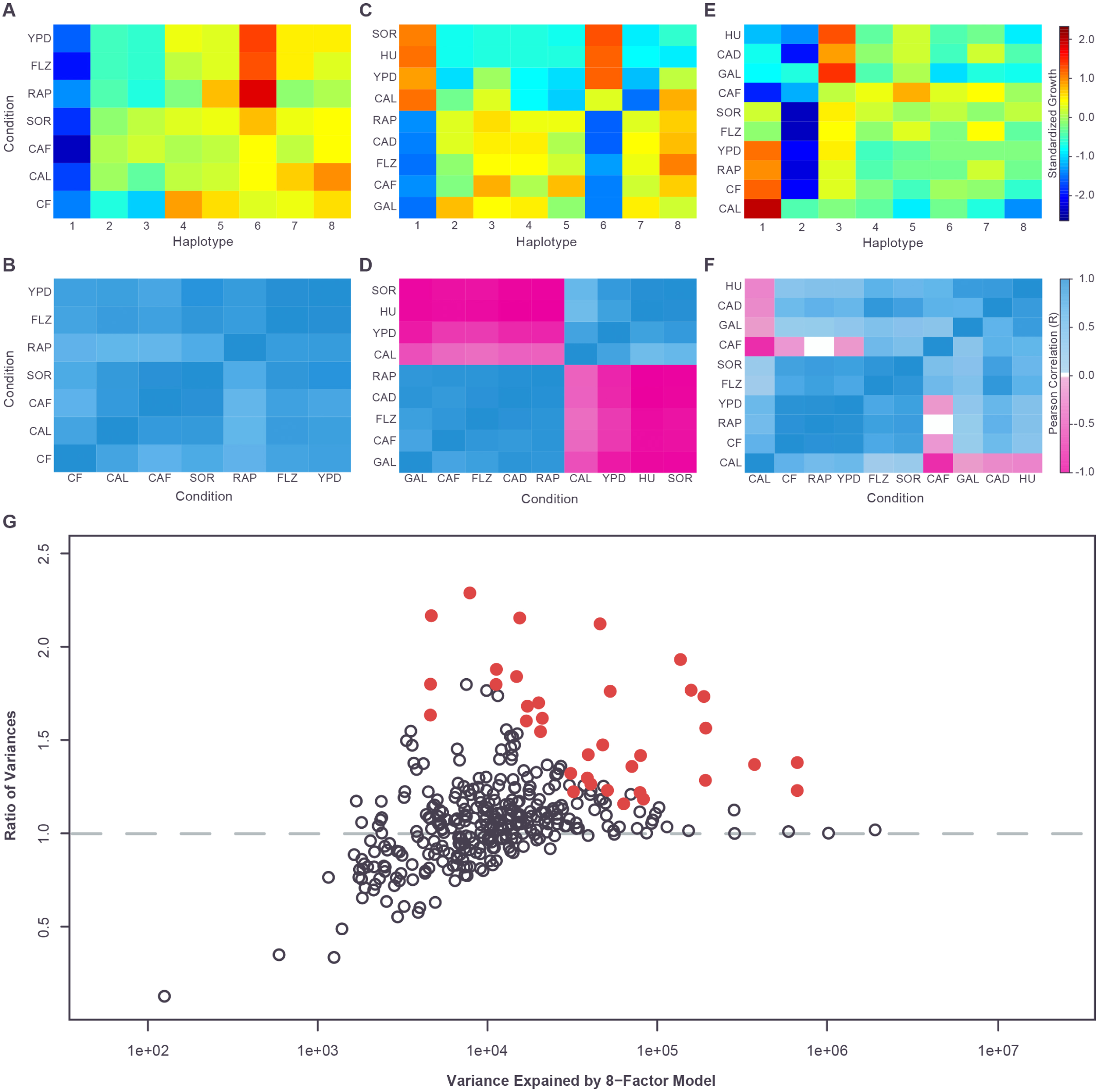
Genetic heterogeneity of QTL and pleiotropic QTL behavior. Standardized haplotype effect sizes (relative to population mean) at *HPF1* (**A**), *WHI2* (**C**) and *FLO11* (**E**). Haplotype effect size correlations between conditions for *HPF1* (**B**), *WHI2* (**D**) and *FLO11* (**F**). (**G**) Ratio of variance explained by full 8-haplotype model versus binarized model, estimated by cross-validation, for all QTL in additive models for all conditions at 72h. Loci demonstrating significantly (p<0.05 after Holm correction) better performance of the 8-haplotype model (i.e. polymodal loci) are indicated in red.

A more complex pattern of correlations was seen at *FLO11* (Table S9: https://doi.org/10.5281/zenodo.17362654). At this locus, the variation in effect profiles is driven largely by differing behaviors of haplotypes 1, 3, and 6 (Figs 3E and 3F). The effect of haplotype 1 varies from promoting growth in some conditions, through little effect on growth, to reducing growth (relative to the population mean). Haplotype 3 varies from little effect on growth to reducing growth, while haplotype 6 varies from little effect on growth to promoting growth. The combinations of these effects result in varying degrees of similarity between condition profiles, ranging from the nearly identical profiles seen in CF, RAP and YPD to the strong negative correlation seen between the CAF and CAL conditions. (Figs 3E and 3F). These results are consistent with multiple variants at *FLO11* in our mapping population, distributed differently across the eight founder haplotypes and each impacting the growth conditions in distinct ways, i.e. closely-linked context-dependent causative variants. *FLO11* is known to have a large and complex promoter, bound by a number of transcription factors. Variation at these transcription factor binding sites would be expected to show pleiotropic effects if *FLO11* expression is more strongly impacted by certain transcription factors under certain growth conditions relative to others.

Taken together, these results indicate that a small number of highly pleiotropic loci are observed across the set of uncorrelated phenotypes we examined, but the allele/haplotype effects at each pleiotropic locus can be highly variable across conditions/phenotypes. Multiple closely-linked variants may contribute to this heterogeneity, with effects on phenotype that vary across the ten conditions. Because these closely linked variants would have different levels of fitness across different phenotypes/conditions, evolutionary selection on this kind of locus may be complex, with tradeoffs between the different phenotype/condition-specific fitness effects.

### QTL often reflect multiple, closely linked causative variants

Our pleiotropy analysis suggested that multiple closely linked variants at a single gene could contribute to phenotypic variance across different conditions. We next set out to investigate whether any of the single-condition QTL in our dataset also result from the effect of multiple, closely linked variants rather than the impact of a single variant. The phenomenon where multiple variants at a single locus contribute to phenotypic variation is described as allelic heterogeneity. Even with our highly powered resource, it is not feasible to comprehensively map causative variants to the nucleotide level. We therefore instead searched for statistical signals that are inconsistent with a single variant underlying a QTL. In the mapping population, recombination occurs only rarely between closely-linked variants, with∼0.09 crossover per 10 kb after 3 rounds of meiosis (described in the companion paper: https://doi.org/10.1101/2025.10.15.682626), so that the parental haplotypes remain mostly intact over distances of <10 kb. This means that the effect of closely-linked variants is difficult to isolate through recombination. However, unlike a pairwise cross where there are only two haplotypes, in a multi-parent cross, there can be as many haplotypes as there are founder strains, which in the case of CYClones is up to eight distinct multi-locus haplotypes. In the case of a single causative (biallelic) variant underlying a QTL, each of these eight haplotypes will always be associated with one of two phenotypic states (bimodality) (Fig S2A), depending on the allele that is present. In the case of multiple-closely linked variants, bimodality can still occur, but the eight haplotypes can also be associated with more than two phenotypic states (polymodality) through imperfect linkage disequilibrium of the variants among the eight founder strains (Fig S2B). Therefore, although genetic heterogeneity can also underlie a bimodal distribution, any QTL in CYClones with a polymodal haplotype phenotype pattern is inconsistent with a single causative variant.

On this basis, we examined each QTL identified in the additive models for each of our ten conditions and, using cross validation and controlling for the effect of all other QTL, compared the variance explained by fitting phenotype effects for all eight haplotypes, to that explained by using a binary haplotype recoding. The binary recoding was chosen using k-means clustering (k=2) of the estimated phenotype effects of the eight haplotypes (Materials and Methods). Our results showed that many QTL, including several of the strongest, are essentially bimodal, with little difference between the variance explained by the full eight haplotypes versus the binary recoding (Fig 3G and Table S10: https://doi.org/10.5281/zenodo.17362654). In contrast, other strong QTL showed a substantial and significant (p-value<0.05 after Holm correction, see Materials and Methods) increase in the variance explained by the eight-haplotype model versus the binary model, consistent with polymodality (Fig 3G and Table S10: https://doi.org/10.5281/zenodo.17362654). In particular, the pleiotropic *FLO11* locus displayed strong polymodality across several growth conditions, consistent with the idea that multiple closely linked variants are responsible for the differences in *FLO11* allele effect patterns observed between conditions. In total 36/349 (10.3%) of QTL were significantly polymodal and, therefore, QTL arising from the effect of multiple linked loci are relatively common in our dataset. In addition, the polymodal loci represent only a minimum estimate of the number of QTL that result from allelic heterogeneity. As mentioned above, although QTL with polymodal effect patterns can only arise from multiple closely-linked causative variants, closely-linked variants can also produce QTL with bimodal effect patterns. This can occur if the variants have indistinguishable effect sizes (Fig S2C), or if there are only two allele patterns present among the founder haplotypes (Fig S2D).

### Allelic heterogeneity and introgressions define a complex set of non-additive genetic interactions in the galactose pathway

To investigate the statistical signals generated by our QTL analysis in the context of a known biological process, we compared our growth results from the galactose condition to the composition and structure of the well-characterized galactose utilization pathway. The enzymatic components of this pathway consist of the Gal2 galactose transporter and the catalytic enzymes Gal1, Gal7, Gal10, Pgm1, and Pgm2 (Fig 4A). While *GAL1*, *GAL7* and *GAL10* (hereafter *GAL1/7/10*) are a cluster of adjacent genes on chromosome II, the remaining genes are dispersed across the genome. The pathway is transcriptionally regulated by the Gal4 transcriptional activator, whose activity is modulated by Gal3 and Gal80, forming a three-protein regulatory circuit (Fig 4A).

**Fig 4.**
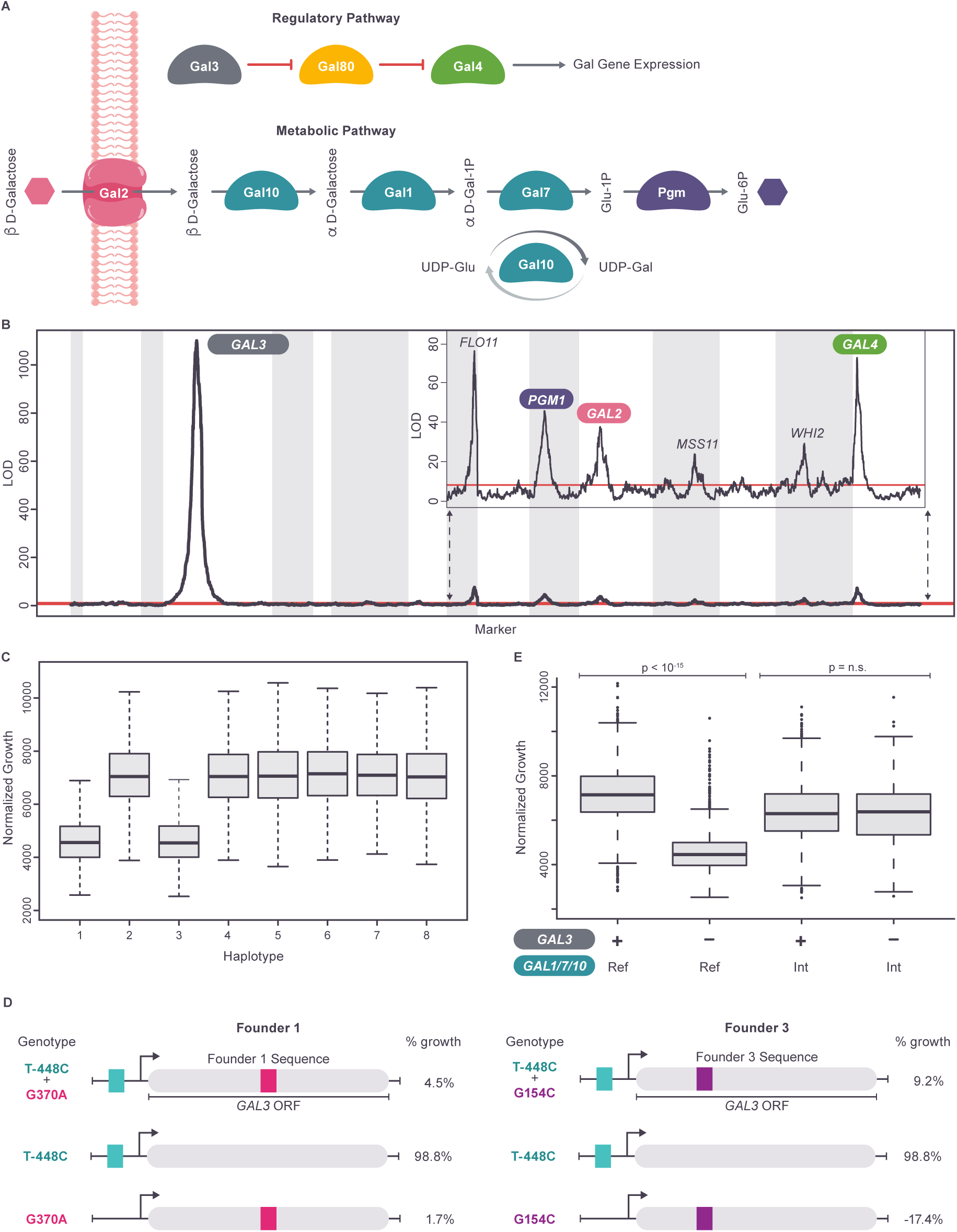
Effect of GAL3 locus on growth on galactose. (**A**) Yeast galactose utilization pathway with genes colored by locus (*GAL1/7/10* form a single locus). (**B**) LOD scores for linkage mapping on galactose. Strong peaks at GAL pathway genes indicated. The *FLO11*, *WHI2* and *MSS11* QTL are not specific to the GAL condition. Insert rescales Y axis to highlight strong QTL other than *GAL3.* 5% family-wise significance threshold in red. Chromosomes alternating in grey and white. (**C**) Growth on galactose for strain subpopulations defined by *GAL3* haplotype. (**D**) Effect on founder 1 and founder 3 growth on galactose after transformation with plasmids expressing their own *GAL3* allele (top) or derivatives (below). Growth is scaled relative to transformation with the S288c reference allele of *GAL3* (100%) and an empty vector (0%). (**E**) Masking epistasis between the *GAL1/7/10* introgression and *GAL3* haplotypes, classified as loss-of-function (*GAL3*-, haplotypes 1&3) or functional (*GAL3*+, remaining haplotypes).

In our dataset, the genetic architecture of the GAL condition is dominated by a single QTL (Fig 4B), which mapped to single gene resolution at *GAL3* and explained 42% of the total variance and 72% of narrow sense heritability (Fig 4B and Tables S3, S8: https://doi.org/10.5281/zenodo.17362654). The phenotype distributions of the eight *GAL3* haplotype subpopulations are bimodal (Fig 4C), with haplotypes from founders 1 (Tuba wine, Philippines) and 3 (Raffia wine, Nigeria) associated with low growth (*GAL3^-^*) and the remaining six haplotypes associated with high growth (*GAL3^+^*). Because these results are consistent with a single bi-allelic causative locus, we examined the *GAL3* sequences of the 8 founder strains and identified a single shared SNV (T-448C) in the *GAL3* promoter of founders 1 and 3. To resolve the *GAL3* locus to the quantitative trait nucleotide (QTN), we therefore constructed and assayed substitution mutations that changed this SNV in the promoters of founders of 1 and 3 to the reference genome allele (found in the other six haplotypes). Surprisingly, reverting this shared allele (T-448C) had no effect on the galactose growth of these founder strains (Fig 4D). Instead, we discovered that two different, closely linked, missense alleles in the coding regions of *GAL3,* one unique to founder 1 (c.G370A, p.Gly124Arg) and another unique to founder 3 (G154C/Gly52Arg), were sufficient to confer the loss-of-function phenotype (Fig 4D). That is, the *GAL3* QTL results from two different *GAL3* variants producing the same low-growth phenotype (allelic heterogeneity) rather than from the effect of a single biallelic locus. This is an example of allelic heterogeneity existing even when the phenotypes associated with the eight founder haplotypes are almost perfectly bimodal (see preceding section and Table S10: https://doi.org/10.5281/zenodo.17362654).

Because genetic variation at *GAL3* has such a strong effect on the GAL phenotype, non-additive pairwise interactions involving *GAL3* also have the potential to significantly impact the phenotype. We therefore scanned for any significant non-additive pairwise interactions between the *GAL3* locus and each of the scaffolding markers. This identified a highly significant interaction between *GAL3* and the *GAL1/7/10* region (Fig S3), despite there being only a very weak QTL peak detected at the *GAL1/7/10* region alone in the original single marker QTL scan (Table S3: https://doi.org/10.5281/zenodo.17362654), i.e. *GAL1/7/10* behaves as a *GAL3* modifier locus. Characterizing this interaction in more detail, we observed that in strains having haplotype 6 at the *GAL1/7/10* locus, variation at the major *GAL3* locus had no effect on growth, with *GAL3^+^* and *GAL3^-^* strains displaying the same mean phenotypes (Fig 4E), i.e. full masking epistasis. This striking result underscores that even a major-effect locus like *GAL3* can exhibit strong context dependence, with its phenotypic consequences entirely contingent on variation at interacting loci such as *GAL1/7/10*.

Founder strain 6 (South African soil) has previously described introgressions of non-*cerevisiae* sequences at the *GAL1/7/10* locus (Fig S4A), as well as at *GAL2* and *PGM1* [27]. Our study therefore identifies (p-value=1.1x10^-149^) a strong non-additive interaction between the presence or absence of the introgression at *GAL1/7/10* and variation at *GAL3*. In contrast, no significant pairwise interaction was observed between *GAL3* and the introgressions at *GAL2* (p=0.81) and *PGM1* (p=0.13). A previous study [16] determined that expression of the introgressed genes (in founder 6) is not repressed in glucose, suggesting their expression may have escaped downregulation by the Gal3-Gal80-Gal4 regulatory circuit. Our *GAL3*-introgression interaction results suggest that it is specifically the *GAL1*/7/10 introgression that is responsible for the escape from control by the regulatory circuit.

In addition to these results, our data confirm [27] that strains harboring introgressions at all three loci grow significantly better on galactose than strains with non-introgressed (“native”, i.e. haplotypes 1:5,7,8) alleles (one-sided t-test, p=3.8x10^-3^) and that combinations of native *PGM1* and the introgressed alleles of *GAL1/7/10* and *GAL2* exhibit significantly lower growth (one-sided t-test, p=1.0x10^-19^) than strains harboring only native alleles (Fig S4B). This low growth is likely due to the build-up of toxic intermediates caused by combining high-activity introgressed alleles and low activity native alleles at these three loci. Our results extend the previously observed 3-way genetic interaction between introgressed and native alleles at *PGM1*, *GAL1/7/10* and *GAL2* [27] into a significant (p=7.5x10^-171^) 4-way interaction that includes *GAL3* (*GAL3^+^* vs *GAL3^-^*). Altogether, these findings underscore the context dependence of genetic effects in the galactose utilization pathway, as the functional impact of specific variants is modulated by the surrounding genetic background, including variation in both the regulatory and structural components of the GAL pathway. In particular, introgressed alleles may alter or bypass native regulatory mechanisms, enabling components of the pathway to escape canonical transcriptional control and thereby reshaping the genetic context in which variants in regulatory genes, such as those at *GAL3*, exert their effects.

### Allele dependent stratification of the galactose pathway: subgroups characterized by distinct regulatory states and genetic architectures

The interaction between *GAL1/7/10* and *GAL3* stratifies the mapping population into subgroups of strains in three different regulatory states. In strains with the *GAL1/7/10* introgression, the effect of variation at the master *GAL3* regulator is suppressed, and variation in growth on galactose is dominated by the interactions between the introgressed and native alleles at *PGM1* and *GAL2* (Figs 5A and S4B and Table S11: https://doi.org/10.5281/zenodo.17362654). Conversely, in strains without the *GAL1/7/10* introgression, growth is dominated by variation at *GAL3*, with the *GAL3^+^* strains inducing the GAL pathway and promoting growth, relative to the *GAL3^-^*strains (Fig S5). To determine the effect of these two regulatory states on genetic architecture, we therefore carried out QTL mapping in the *GAL3^+^*and *GAL3^-^* subgroups (after removing strains with the *GAL1/7/10* introgression). For clarity, we refer to these sets of strains as ‘subgroups,’ to distinguish them from genetically diverged subpopulations. This approach allows us to assess how the phenotypic effects of genetic variants are shaped by context dependence, that is, how their impact varies depending on the regulatory background established by *GAL3* and *GAL1/7/10*.

**Fig 5.**
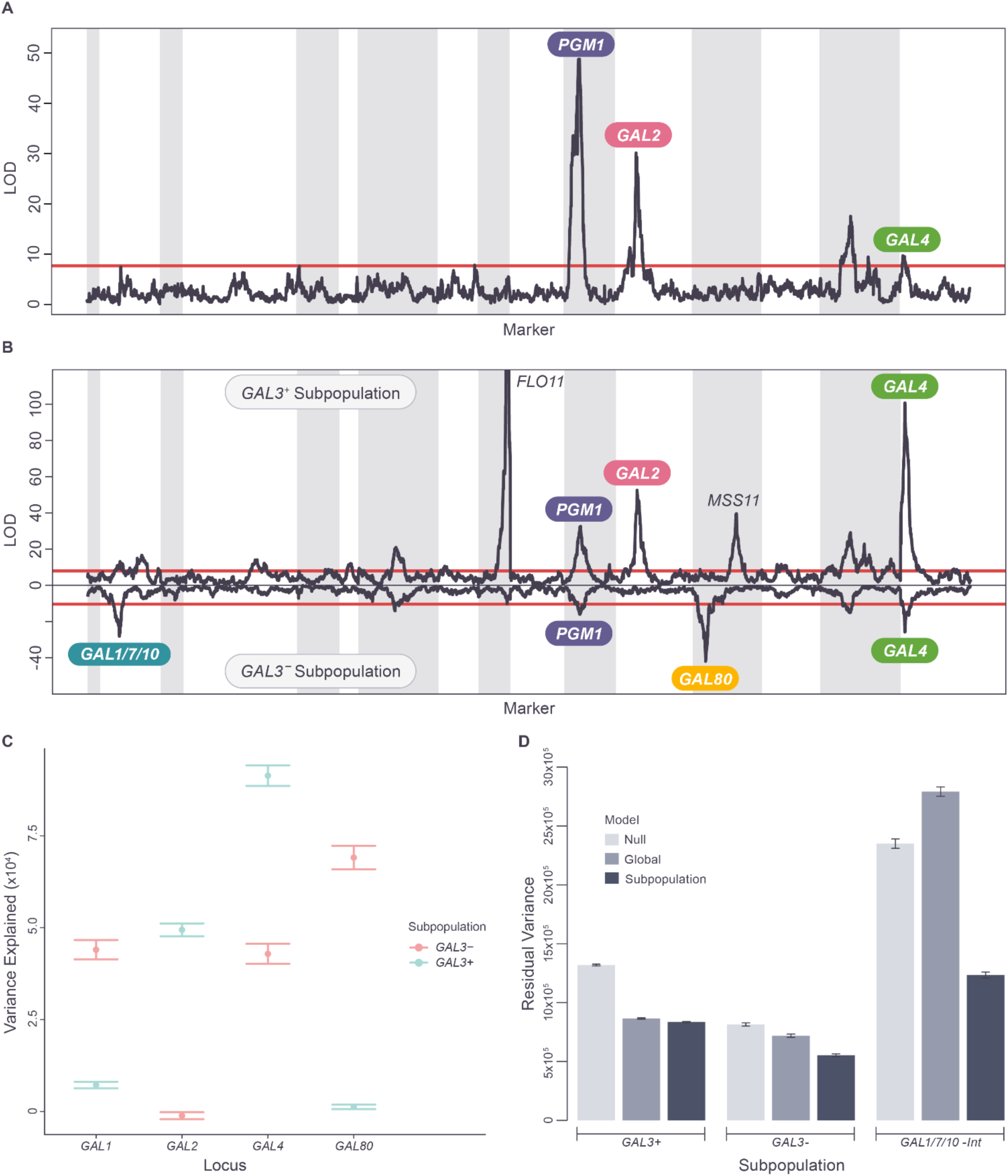
Genetic architecture of growth on galactose. GAL pathway genes labelled when falling in QTL peak 2-LOD confidence intervals, 1% significance thresholds in red. (**A**) QTL mapping in strains with the introgressed *GAL1/7/10* locus. (**B**) QTL mapping in subpopulations lacking the introgressed *GAL1/7/10* locus, split based on *GAL3*+ vs *GAL3*-. *GAL3*-LOD scores plotted on negative scale. The *FLO11*, *WHI2* and *MSS11* QTL are not specific to the GAL condition. (**C)** Variance (+/- standard error) explained by individual GAL loci in each subpopulation (in the absence of the *GAL1/7/10* introgression) using random sampling cross validation. (**D**) Performance of global and subpopulation-specific additive models on each subpopulation, assessed by random sub-sampling validation (mean residual estimate +/- SE values shown).

The QTL landscapes of the two subgroups displayed substantial differences (Fig 5B and Tables S11, S12 and S13: https://doi.org/10.5281/zenodo.17362654). First, the effect of variation at QTL containing the regulatory genes *GAL80* and *GAL4* differed between the two subgroups in a manner consistent with the logic of the regulatory circuit (Fig 4A). A significant QTL was seen at the *GAL80* locus (mapped to a 5 kb interval) in the *GAL3^-^* strains, while no significant QTL was observed at *GAL80* in the *GAL3^+^* strains (Figs 5B and 5C and Tables S11, S12 and S13: https://doi.org/10.5281/zenodo.17362654). Genetic variation in *GAL80* only has a significant phenotypic impact in the presence of the loss of function alleles of *GAL3*, when Gal3 no longer inhibits Gal80 (Fig. 4A). Conversely, the phenotypic effect of variation at the *GAL4* locus is significantly stronger (two-sided t-test p=1.2.x10^-26^) in the *GAL3^+^* subgroup than in the *GAL3^-^* subgroup (Figs 5B and 5C and Tables S11, S12 and S13: https://doi.org/10.5281/zenodo.17362654), which is also consistent with the structure of the regulatory circuit (Fig 4A), as Gal4 is derepressed in the *GAL3^+^*strains.

In addition to differences in the phenotypic effects of variation in the *GAL4* and *GAL80* regulatory genes, the *GAL3*⁺ and *GAL3*⁻ subgroups also displayed different effects of variation at loci involved in mechanistic steps of the galactose utilization pathway (Figs 5B and 5C). There is a significant QTL peak at *GAL2* only in the high-growth subgroup (mapped to an 11 kb interval), while the GAL2 locus has no significant effect on phenotype in the low-growth subgroup. Conversely, variation at *GAL1/7/10* (haplotype 6 excluded) had a stronger effect on growth in the low-growth subgroup (two-sided t-test p=2.8.x10^-25^; Figs 5B and 5C). This differential sensitization to variation at *GAL2* vs *GAL1/7/10* may reflect differences in the degree to which the activity of each protein is limiting when the GAL pathway is induced (*GAL3^+^*) vs non-induced (*GAL3^-^*). These results illustrate a strong form of context dependence, in which the genetic effects of both regulatory and structural loci shift depending on the regulatory state imposed by *GAL3* and *GAL1/7/10*. The distinct genetic architectures associated with the three identified subgroups of the mapping population (*GAL1/7/10* introgression, *GAL3^+^*, *GAL3^-^*) mean that a single additive model fitted to the entire population cannot describe the genotype-phenotype relationship of all three subgroups well. In fact, a global model appears to capture the genetic architecture of the *GAL3^+^* subgroup (69% of strains) quite accurately (Fig 5D), reducing residual phenotypic variance significantly relative to a subgroup null model (34% reduction, two-sided t-test p=4.1.x10^-108^), performing similarly to a subgroup-specific additive model (37% reduction relative to null model). The global model performs less well (Fig 5D) on the *GAL3^-^* subgroup (22% of strains), reducing residual phenotypic variance significantly (two-sided t-test p=1.2.x10^-6^) relative to a subgroup null model but performing less well than a subgroup-specific model (12% vs 32% reduction, two-sided t-test p=4.2.x10^-17^). On the *GAL1/7/10* introgressed subgroup (9% of strains), the global model performs very poorly (Fig 5D), actually significantly increasing residual phenotypic variation relative to a subgroup null model (19% increase, two-sided t-test p=2.1.x10^-13^) and performing much worse (two-sided t-test p=4.9.x10^-76^) than a subgroup-specific model (48% decrease in residual variance relative to null).

It is important to note that in the analyses described in this section, the subgroups analyzed share the same patterns of variation at all loci other than the *GAL1/7/10* and *GAL3* regions used to define the subgroups. Therefore, the subgroups are not genetically diverged subpopulations, instead, they display different genetic architectures despite sharing the same genome-wide patterns of genetic variation, a striking example of context-dependent genetic effects driven by allele-specific regulatory states. We refer to this phenomenon, where allelic variation determines genetic <u>architecture</u>, as “allele determined stratification”. In the case of our GAL results, these distinct architectures appear to result from distinct regulatory states associated with allele state at *GAL1/7/10* and *GAL3*.

### QTL for growth on cadmium chloride: an example of quantitative allele dependent stratification

Our analysis of the galactose growth condition revealed that allele-dependent stratification corresponded to qualitatively distinct regulatory states. Analysis of the major QTL for growth on cadmium chloride extended this qualitative model to characterize a quantitative, allele-determined transition between genetic architectures, a striking example of context dependence, where the functional effects of genetic variants shift continuously with changes in the internal cellular environment.

The strongest single QTL identified in our analysis was observed in the cadmium chloride (CAD) condition and maps to the right end of chromosome II (LOD = 2434; 70% of total variance) (Tables S3 and S8: https://doi.org/10.5281/zenodo.17362654). Further analysis determined that Founder 2 (United States soil) contains an introgression replacing several genes in this region with orthologs from another species (Fig S6 and File S1https://doi.org/10.5281/zenodo.17362654). Reads from this introgressed region align poorly to the budding yeast reference sequence, causing undercounting of haplotype 2. Therefore, to improve sensitivity for identifying strains inheriting introgressed sequences from this region, we re- haplotyped the right end of chromosome II using an introgression-aware pipeline (Methods).

After re-haplotyping and performing linkage analysis, the QTL in this region (Table S14: https://doi.org/10.5281/zenodo.17362654) became even stronger (max LOD = 2624; 72% of total variance), dominating resistance to cadmium chloride with an almost Mendelian genetic architecture and mapping to a 2.4 kb interval (Table S15: https://doi.org/10.5281/zenodo.17362654) that includes the gene *PCA1*, which encodes a cadmium efflux pump [28]. The effect of the *PCA1* haplotype on growth in CAD was complex. Strains harboring the introgressed *PCA1* haplotype grow much more rapidly than the other strains, but there are also large differences in growth between the seven non-introgressed haplotypes at *PCA1* (Fig 6A), with haplotype 1 growing best and haplotype 7 growing worst. This strongly suggests that multiple *PCA1* variants are impacting function (allelic heterogeneity), and that these variants are in linkage disequilibrium among the seven (non-introgressed) founder strains.

**Fig 6.**
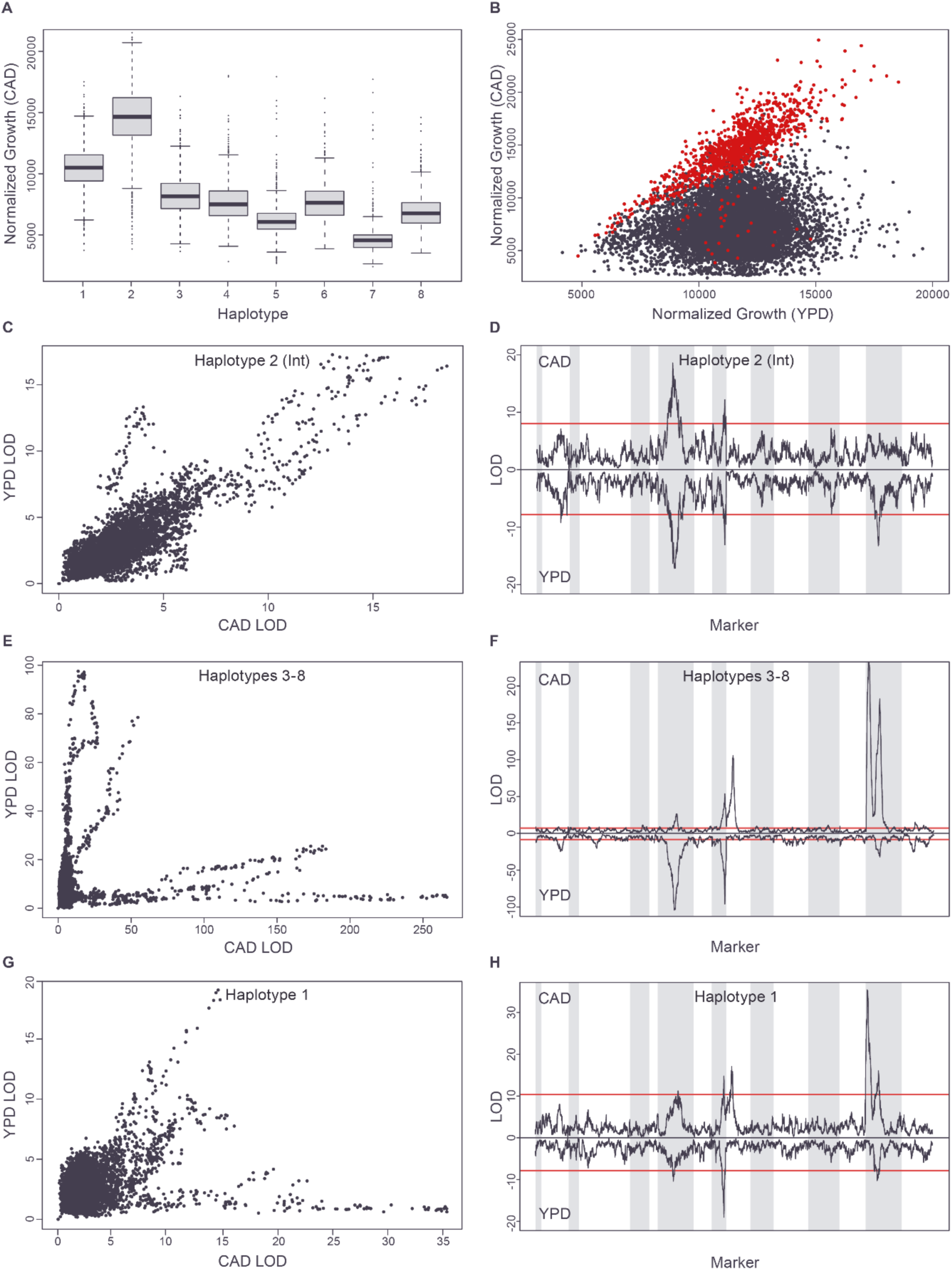
PCA1 haplotypes define a quantitative shift between two genetic architectures on CAD. (**A**) Strain growth on CAD based on *PCA1* haplotype. (**B**) YPD vs CAD growth, strains with the *PCA1* introgression (haplotype 2) in red, rest in black. (**C,E,G**) LOD scores for each scaffolding marker on YPD versus CAD for each of the identified subpopulations defined by *PCA1* haplotype (after accounting for the effect of *PCA1* haplotype for the combined haplotype 3-8 population). (**D,F,H**) LOD scores for each scaffolding marker on CAD and YPD plotted against scaffolding marker number for each of the identified subpopulations defined by *PCA1* haplotype, with YPD LOD scores plotted on a negative scale. Chromosomes indicated as alternating grey and white backgrounds. 1% significance thresholds in red.

For the entire mapping population, growth in the presence (CAD condition) and absence (YPD condition) of cadmium is essentially uncorrelated (R^2^=0.029), although growth on YPD does define an upper boundary for growth on CAD (Fig 6B). However, strains with the introgressed *PCA1* haplotype grow so well on CAD that we hypothesized they might behave as if they were growing in the absence of this heavy metal stressor. Indeed, growth rates on CAD and YPD are strongly and significantly correlated among strains carrying the introgressed *PCA1* gene (R^2^ = 0.61; p-value = 5.9x10^-248^), and these strains fall on the upper boundary of growth on CAD (Fig 6B and Fig S7). In addition, the QTL profile of the introgressed strains on CAD and YPD is almost identical (Figs 6C and 6D and Table S16: https://doi.org/10.5281/zenodo.17362654). These results are consistent with the *PCA1* introgression making strains fully resistant to cadmium, so that their growth is constrained by the same genetics that determine their growth in the absence of cadmium (i.e. on YPD).

In contrast to the introgressed strains, strains having haplotypes 3-8 at the *PCA1* locus show lower levels of resistance to cadmium (Fig 6A) and display low correlation between growth on YPD and CAD (R^2^<0.085 for every haplotype), (Fig S7). Similarly, the genetics determining growth on YPD and CAD differ strongly for these strains (Figs 6E and 6F). On YPD, the QTL trace of the combined haplotype 3-8 strains is very similar to that seen for strains with the *PCA1* introgression (Figs 6D and 6F and Table S16: https://doi.org/10.5281/zenodo.17362654, R^2^=0.64), representing a shared YPD genetic architecture. On CAD, after accounting for the effect of *PCA1*, haplotypes 3-8 share a single genetic architecture (Fig S8) with the first principal component explaining 87% of standardized variance in their LOD profiles. This architecture is very distinct from the common YPD architecture and includes major peaks including the cadmium-related genes *SMF1*, a transporter involved in the transport of cadmium and other divalent ions (mapped to a 3.4 kb interval), and *GSH1*, a glutathione biosynthesis gene involved in the cellular response to cadmium [29] (mapped at single gene resolution to a 1.6 kb interval) (Table S17: https://doi.org/10.5281/zenodo.17362654). Therefore, among strains with these six *PCA1* haplotypes (haplotypes 3-8), the degree of growth on cadmium, after accounting for the effect of the major *PCA1* locus, appears to be largely determined by additional loci specifically affecting cadmium resistance rather than the genetics determining growth on YPD.

The final set of strains, with haplotype 1 at the *PCA1* locus, shows the second highest mean level of growth on cadmium (Fig 6A). Like the introgressed subgroup, these strains display a correlation between growth on cadmium chloride and on YPD, but lower than that seen with the introgressed strains (R^2^=0.28 vs 0.61; Fig S7). On YPD, the haplotype 1 strains display the same genetic architecture seen in the other *PCA1* subgroups (Fig 3H and Table S16: https://doi.org/10.5281/zenodo.17362654), while the QTL landscape of these strains on CAD is a mixture of the cadmium resistance architecture seen with haplotypes 3-8 and the YPD-like architecture seen in the introgressed strains (Fig 6G and 6H and Table S16: https://doi.org/10.5281/zenodo.17362654). These results are consistent with a quantitative genetic transition between growth constraint by cadmium chloride and growth constraint by YPD, a form of context dependence in which the genetic architecture is modulated by the physiological consequences of allelic variation at *PCA1*. Because Pca1 exports cadmium ions from the cell, the CAD-specific and YPD-like constraint regimes associated with the different *PCA1* alleles may reflect varying levels of toxic intracellular cadmium associated with different levels of Pca1 activity. In this model, *PCA1* haplotypes 3-8 result in relatively low levels of Pca1 activity, leading to high levels of toxic intracellular cadmium and a cadmium-specific genetic architecture constraining cell growth. In contrast, *PCA1* haplotype 2 results in relatively high export activity, leading to low levels of intracellular cadmium and cells with a YPD-like genetic architecture constraining their growth. Finally, *PCA1* haplotype 1 leads to intermediate levels of export activity and intermediate levels of intracellular cadmium, placing strains with this allele in the transitional zone between the two constraint regimes and producing a mixed genetic architecture.

## Discussion

The CYClones multiparent mapping population provides a powerful new tool for delineating the genetic architecture of complex traits that can identify even weak effect QTL with high mapping resolution. By combining dense recombination, high-resolution haplotyping, and highly reproducible, high-throughput phenotyping, CYClones enables the identification of large numbers of quantitative trait loci (QTL) across a wide range of biological conditions. Importantly, the mapping resolution is sufficient to localize many of these loci to single genes or small sets of candidate genes, facilitating direct experimental validation and mechanistic insight.

This level of resolution and throughput, rarely achievable in more complex organisms, demonstrates the enduring value of yeast as a model system for uncovering general principles of genetic architecture. It also highlights the utility of model organisms for translating genotype-phenotype associations into causal mechanisms. Ultimately, the power of CYClones lies not only in its ability to detect and localize QTL, but also in its capacity to characterize genetic complexity, including pleiotropy, context-dependent effects, and the contribution of multiple tightly-linked causal variants to phenotypic variation. In this respect, CYClones plays a role in yeast analogous to the Mouse Collaborative Cross in mammals [7]. Both were constructed using a funnel cross design, underscoring how multiparent populations can capture broad genetic diversity and recombination to resolve complex trait architecture across diverse organisms.

### Genetic complexity in the architecture of quantitative traits

One of the most striking features revealed by the CYClones resource is the diversity and richness of genetic architectures underlying quantitative traits. Some traits are dominated by a single, large-effect locus, while others exhibit highly polygenic architectures shaped by many small-effect variants. This variability arises not only from differences in effect size and number of contributing loci, but also from other components of genetic architecture, including pleiotropy, allelic heterogeneity, and epistatic interactions, that modulate phenotypes in condition-specific ways.

Although most loci affected growth in a condition-specific manner, we identified a small number of broadly pleiotropic loci that influenced multiple traits. Notably, *WHI2*, *FLO11*, and *HPF1* each affected growth in at least seven conditions, often with trait-specific effect directions and magnitudes. For instance, at *WHI2* and *FLO11*, the same haplotype promotes growth in one condition while reducing it in another, suggesting that the same functional allele can have opposing fitness consequences depending on cellular context. These findings illustrate that even when pleiotropy is limited to a few loci, it can introduce substantial complexity and asymmetry into genetic architectures. We note that pleiotropy is likely more widespread than what we observed, as we only considered QTL mapped to single-gene resolution. Moreover, the opportunity to identify pleiotropy was limited by the ten specific growth conditions tested, so the QTL we identified may affect many other phenotypes. Indeed, CYClones will become increasingly powerful for understanding genotype–phenotype relationships as additional traits are measured, particularly given that the mapping population and all sequencing and genotyping data are broadly available to the research community.

Our analysis also reveals that allelic heterogeneity is common, with many QTL best explained by the combined effects of multiple, tightly linked causal variants. Across the genome, 10.3% of QTL exhibit significantly polymodal haplotype effect patterns, inconsistent with a single biallelic variant and indicative of multiple functional alleles contributing distinct phenotypic effects. In the galactose condition, we demonstrate this directly at the *GAL3* locus, where two distinct missense mutations in different founder backgrounds independently produce a loss-of-function phenotype. Moreover, *GAL3* interacts epistatically with introgressed alleles at the *GAL1/7/10* locus, and these interactions, along with additional non-additive effects involving introgressions at *GAL2* and *PGM1*, form a complex, multi-locus regulatory network. The phenotypic consequences of these interactions depend on regulatory state and specific combinations of introgressed and native alleles, illustrating how both allelic heterogeneity and epistasis can structure genetic architecture at the pathway level.

Introgression of alleles from other yeast species is a key contributor to the complexity of genetic architectures that we observed. In both the cadmium (CAD) and galactose (GAL) conditions, introgressed alleles reshape cellular physiology in ways that radically alter the genetic architecture of the trait. These introgressed sequences do not act in isolation but interact with native loci, generating condition-specific architectures that defy simplistic additive models. As such, introgression not only contributes to phenotypic variance but also increases architectural diversity across traits. These results provide important context for evaluating the molecular consequences of introgression in more complicated organisms, including Neanderthal and Denisovan introgressed sequences in modern humans [30, 31].

More broadly, these findings emphasize that genetic complexity is not merely a matter of polygenicity. Instead, it encompasses multiple overlapping phenomena, including pleiotropy, allelic heterogeneity, introgression, and context-dependent interactions, that interact to produce rich, condition-specific genotype-phenotype maps. These interlocking layers of complexity highlight the need to understand genetic architecture not just as a static map of effect sizes, but as a dynamic system shaped by regulatory and physiological state.

### Allele-dependent stratification: a distinct form of context dependence

Beyond identifying individual loci and their interactions, CYClones allowed us to define and explore allele-dependent stratification: a striking and underappreciated form of context dependence in which specific alleles at key regulatory or structural loci effectively partition the population into subgroups that experience distinct genotype-phenotype relationships. This phenomenon differs from the traditional notion of “background effects,” such as those commonly described in mouse models, where allelic effects shift subtly depending on broader genomic context. Instead, allele-dependent stratification arises when specific alleles of a small number of genes produce discrete regulatory or physiological states that, in turn, reshape the genetic architecture of the phenotype itself.

In the GAL condition, the *GAL3* and *GAL1/7/10* loci define three such regulatory subgroups, each with distinct genetic architectures. Similarly, in the CAD condition, variation at the gene encoding the Pca1 efflux pump modulates intracellular cadmium levels, effectively stratifying the population into cells that differ in their physiological exposure to cadmium stress. These differences then determine whether genetic variation influencing detoxification pathways or basal growth rate becomes phenotypically relevant. Crucially, in both examples, the genome-wide patterns of variation are shared across subgroups; only the alleles at a small number of loci differ. The resulting architectural differences are therefore not a consequence of population structure or long-term divergence but rather emerge from functionally mediated, allele-specific shifts in internal state, challenging the standard assumption that a single additive model can capture trait variance across a population and underscoring the need for approaches that can incorporate cellular and regulatory state into predictive frameworks.

When faced with such allele-dependent stratification, our results from the GAL condition demonstrate that global additive models default to accurately reflecting the genetic architecture of the largest subgroups while performing poorly on smaller subgroups, whose distinct genetic architectures are not captured in the model. This behavior also explains why allowing non-additive interactions within whole-population models provides little improvement in global performance. Because the model is dominated by the majority subgroup’s architecture, adding interaction terms cannot recover the fundamentally different genetic architectures of the minority subgroups.

While one could in principle attempt to comprehensively stratify a population based on allelic variation at each QTL to identify subgroups with distinct architectures, such exhaustive approaches suffer from the same statistical limitations, namely, power loss and multiple-testing burden, that constrain genome-wide interaction analyses. However, orthogonal data types, such as gene expression, chromatin accessibility, or protein abundance, can offer independent and biologically meaningful insights into cellular state. These data can help guide the identification of relevant subgroups, reducing the search space and enabling targeted modeling of genotype-phenotype relationships within contextually defined strata.

### Implications for human genetic studies

Our findings suggest that allele-dependent stratification may be a general feature of complex trait architecture, with important implications for human genetics and precision medicine. In particular, the presence of a strong disease-risk or protective allele may place individuals into distinct regulatory or physiological states that fundamentally alter the genetic architecture influencing phenotype. For example, individuals carrying loss of function alleles in *CFTR* (leading to cystic fibrosis) or in *PCSK9* (reducing LDL cholesterol and protecting against heart disease) [32] may exhibit different sets of modifying loci than those relevant to individuals without the major effect variant. That is, the genetic modifiers of phenotype may differ depending on whether an individual carries a major-effect allele, such that global polygenic risk models may systematically misestimate risk for subgroups carrying such alleles. While our yeast studies provide proof-of-principle for this phenomenon, these human analogs represent testable hypotheses that warrant investigation.

This principle may extend beyond specific monogenic variants with extreme phenotypes to individuals who fall at the extremes of polygenic trait distributions, those with exceptionally high or low liability scores. At the tails, the physiological or regulatory landscape may be sufficiently perturbed that the relevant genetic modifiers differ from those shaping phenotypes in the center of the distribution. In this way, polygenic extremity itself may define one or more stratified subgroups, with distinct genetic architectures. As a consequence, polygenic risk scores may be particularly prone to misestimating risk in individuals at the extremes, where the underlying assumptions of a uniform genetic architecture no longer hold. Identifying such allele- or state-defined subgroups, through orthogonal data such as transcriptomic or proteomic profiles, could greatly improve the precision and interpretability of risk modeling efforts.

## Author Contributions

Experiments were designed by A.M.D., J.M.A., G.A.C., L.A. and R.S.L. Experiments were performed by L. A., R.S.L., T.S.M., M.S.T. and J.A. under the supervision of A.M.D. Analyses were conducted by G.A.C., under the supervision of A.M.D. and J.M.A. Software was written by G.A.C. The manuscript was written by G.A.C., R.S.L., J.M.A. and A.M.D. and incorporates comments by all other authors.

## Data availability

Strain phenotype and haplotype data are provided as Tables S2 and S6. An R script (File S2) reproduces all analyses described in the Results. All supporting datasets are publicly available as supplementary files on Zenodo at https://doi.org/10.5281/zenodo.17362654 (CC BY 4.0 license). PyPlate is available at https://github.com/lacyk3/pnri-projects/tree/Image-Analysis-Demos/Funnel%20Cross%20Project.

## Acknowledgements

We thank Drs. Feng-Yan Bai and Justin fay for providing yeast strains. This work was funded by an NIH/ NIGMS award (R01 GM117119) to A.M.D. and J.M.A., institutional funding from PNRI to A.M.D., and NIGMS award (R01 GM110068) to J.M.A.

---

**Fig S1.**
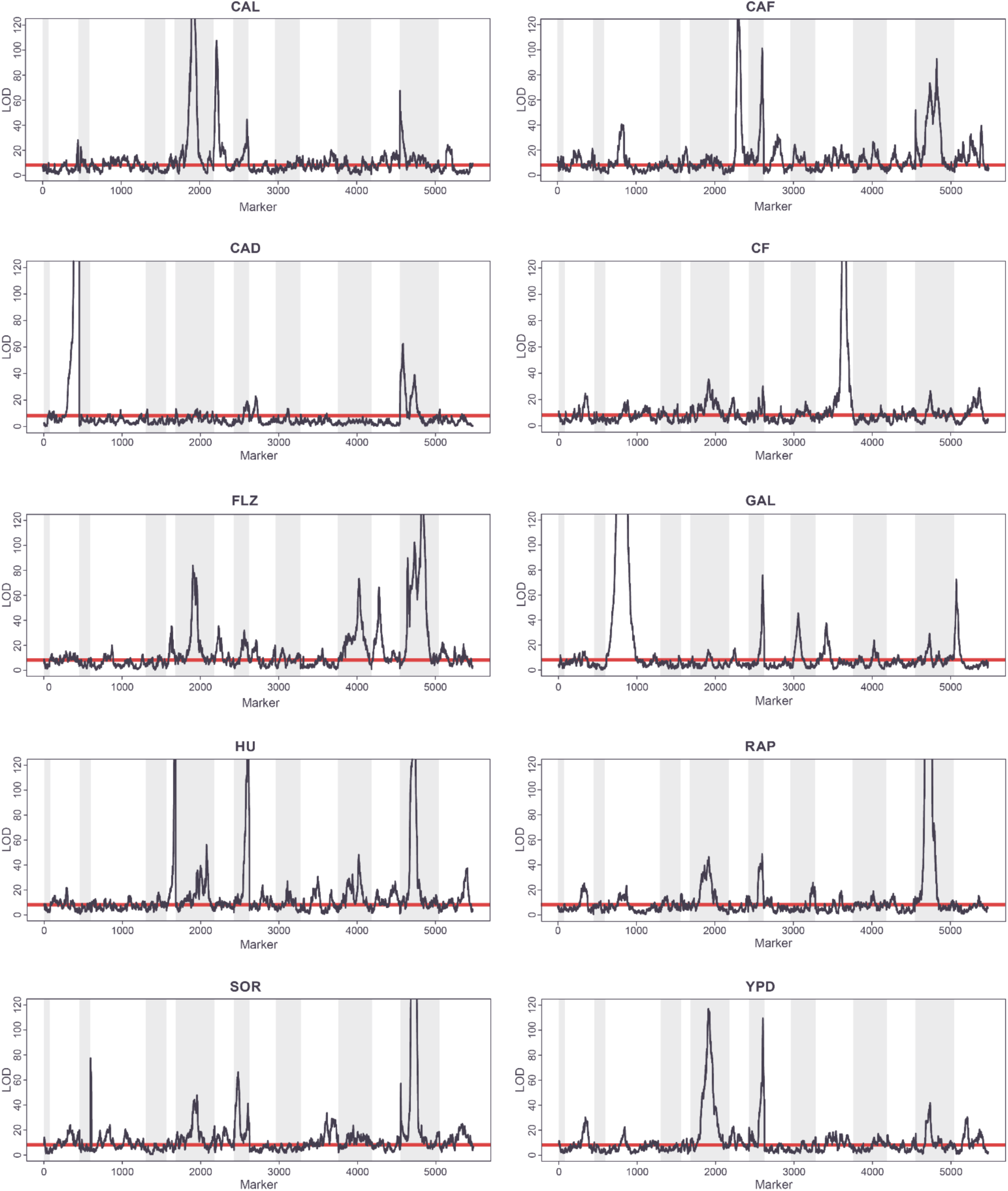
LOD traces for all conditions at 72h. Scaffolding marker set used. Chromosomes indicated by alternating grey and white backgrounds. Red line represents 5% family-wise significance threshold.

**Fig S2.**
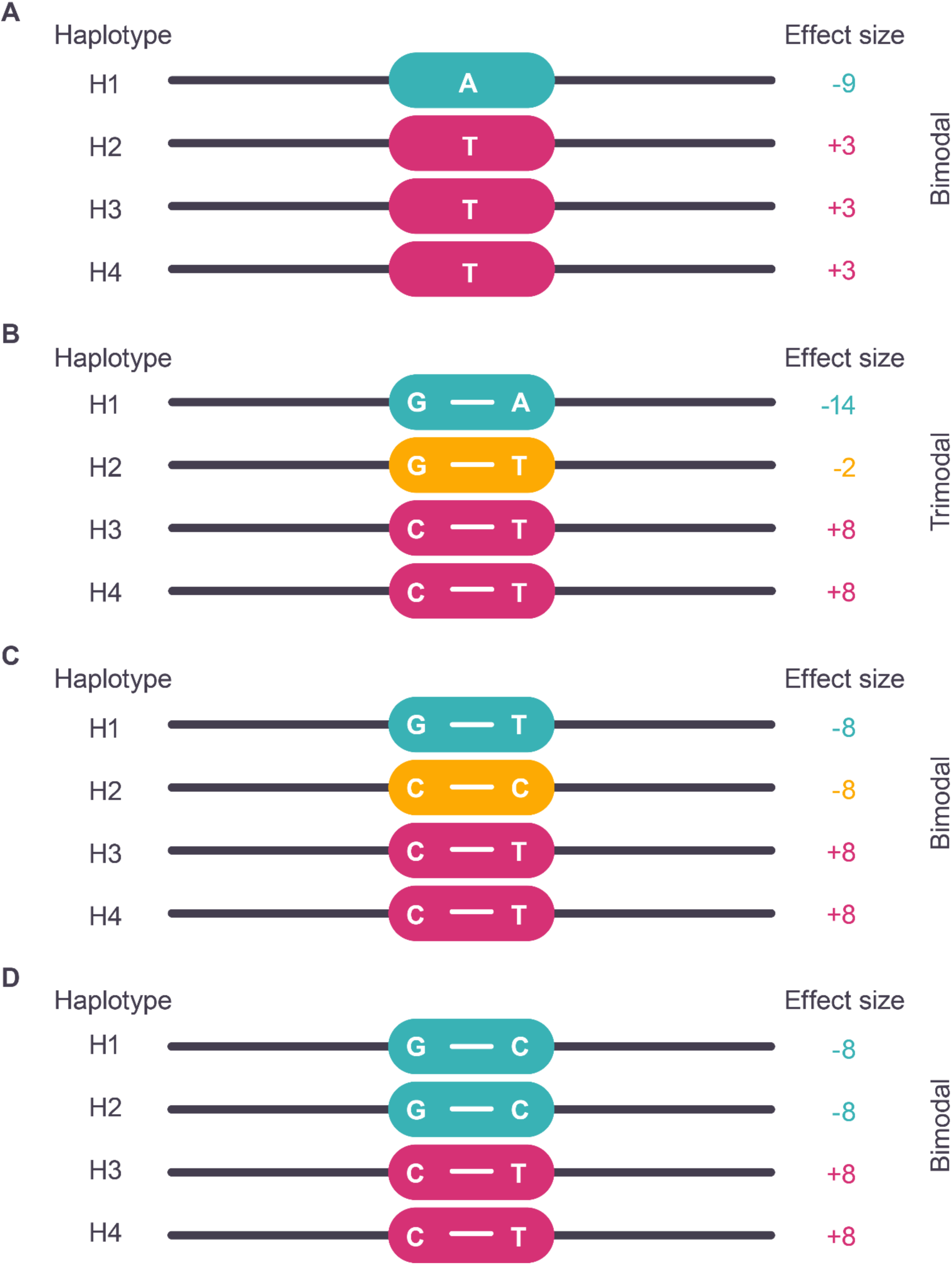
Patterns of haplotype effect sizes at loci with single causative variants or closely linked pairs of causative variants. (**A**) A single causative variant site always produces a bimodal effect distribution. (**B**) Two closely linked causative variant sites permuted among founder haplotypes can produce a polymodal haplotype effect distribution (e.g. G=-2, C=+8, A=-12, T=0). (**C**) Two closely linked causative variant sites permuted among founder haplotypes can produce a bimodal haplotype effect distribution if multiple allele permutations share the same effect size. (**D**) Two closely linked variant sites that are not permuted among founder haplotypes produce a bimodal effect distribution.

**Fig S3.**
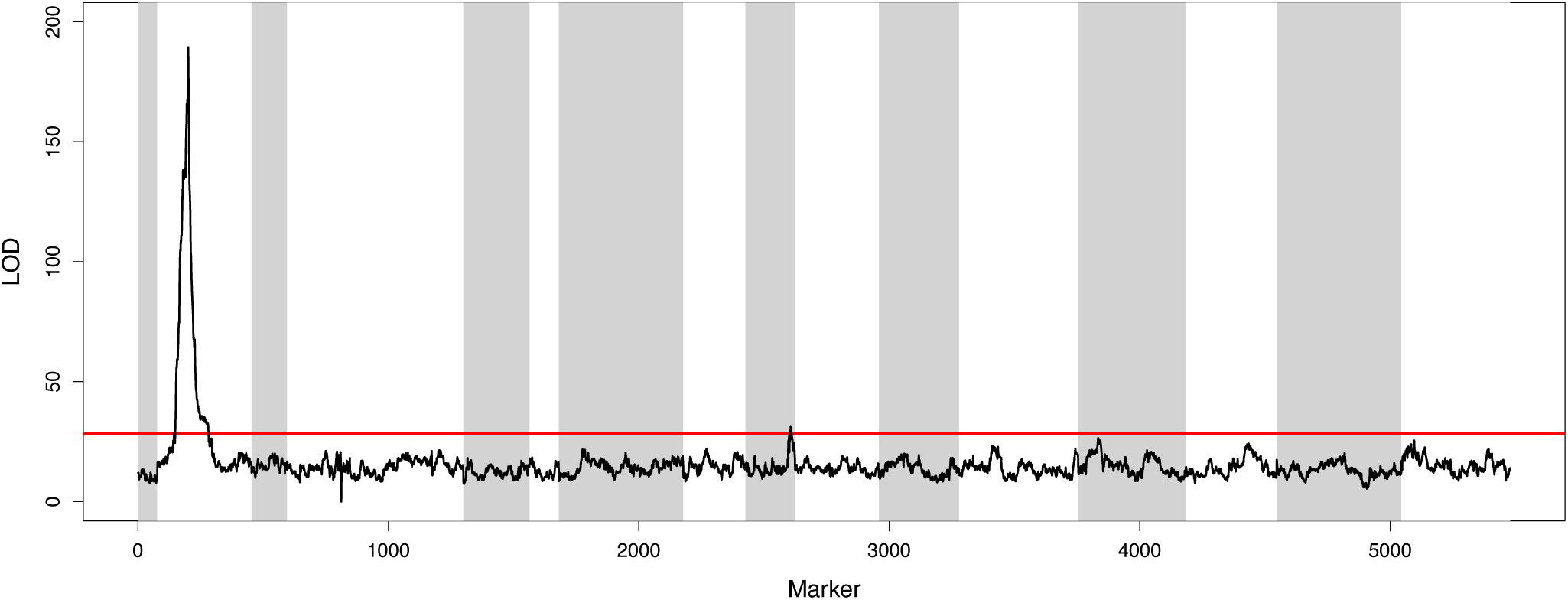
Interaction vs additive LOD trace for *GAL3* maximum marker (chr04_465458) vs all scaffolding markers identifies a strong interaction between *GAL3* and *GAL1/7/10*. Chromosomes indicated by alternating grey and white backgrounds. Red line represents 1% FDR threshold for full GAL interaction vs additive dataset.

**Fig S4.**
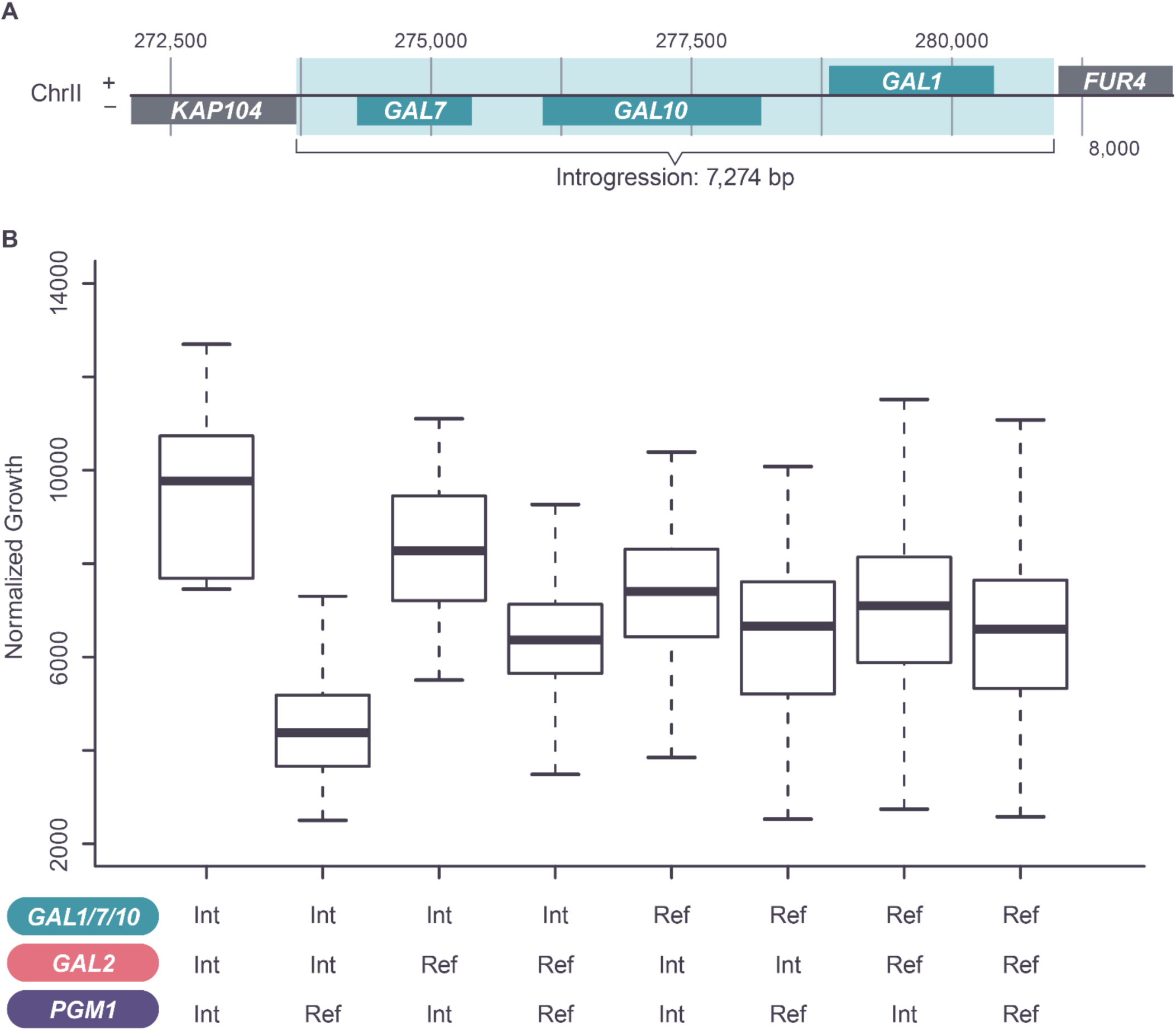
Interactions between reference and introgressed GAL pathway genes. (**A**) An introgression replaces the *GAL7*, *GAL10* and *GAL1* genes with non-native versions in founder 6. (**B**) Effect on strain growth of combining introgressed and non-introgressed (reference) alleles of GAL pathway genes.

**Fig S5.**
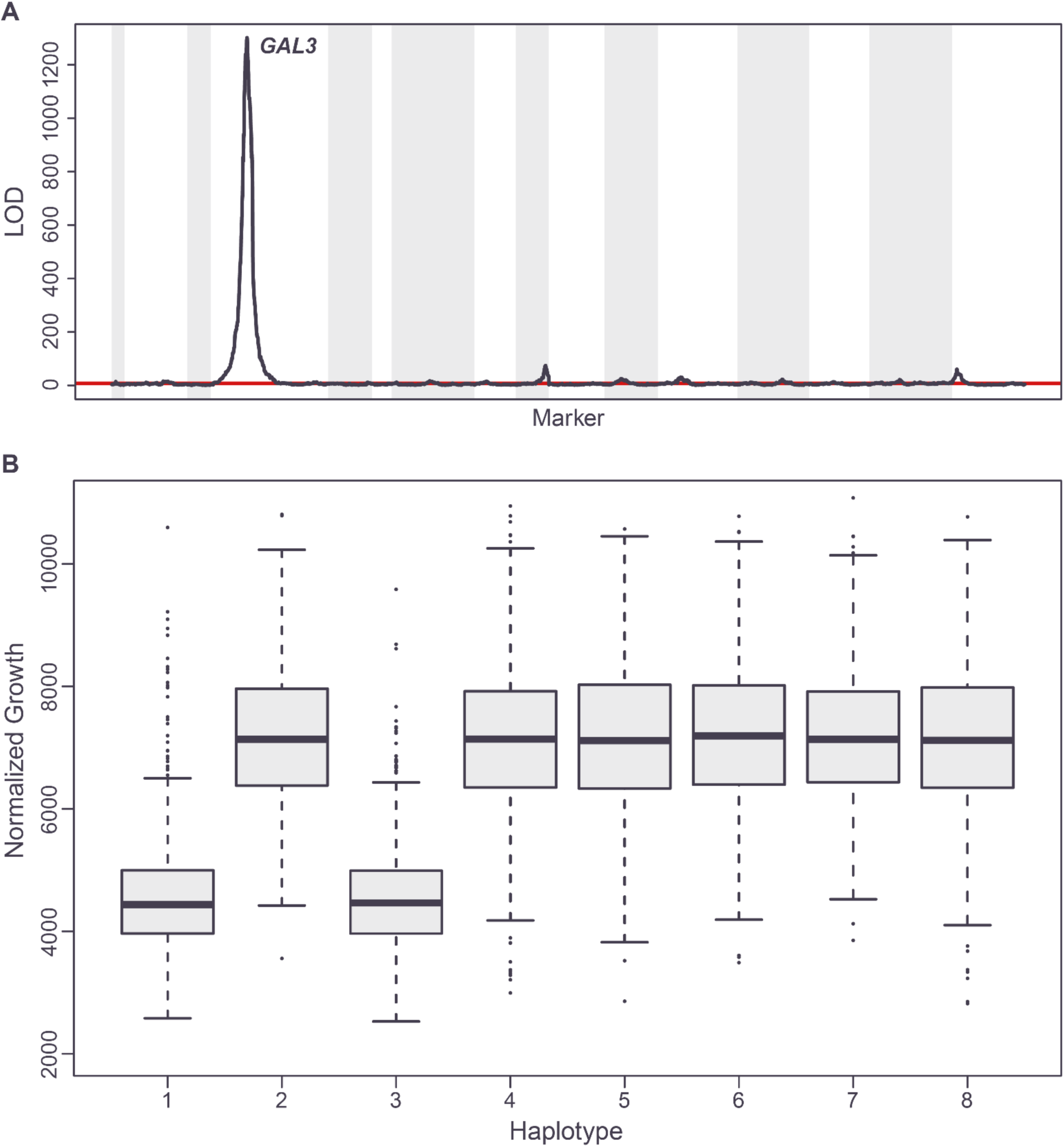
Effect of *GAL3* locus on growth on galactose for strains lacking the *GAL1/7/10* introgression. (**A**) LOD scores for linkage mapping on galactose. Chromosomes alternating in grey and white. 1% significance threshold in red. (**B**) Growth on galactose for strain subpopulations defined by *GAL3* haplotype.

**Fig S6.**
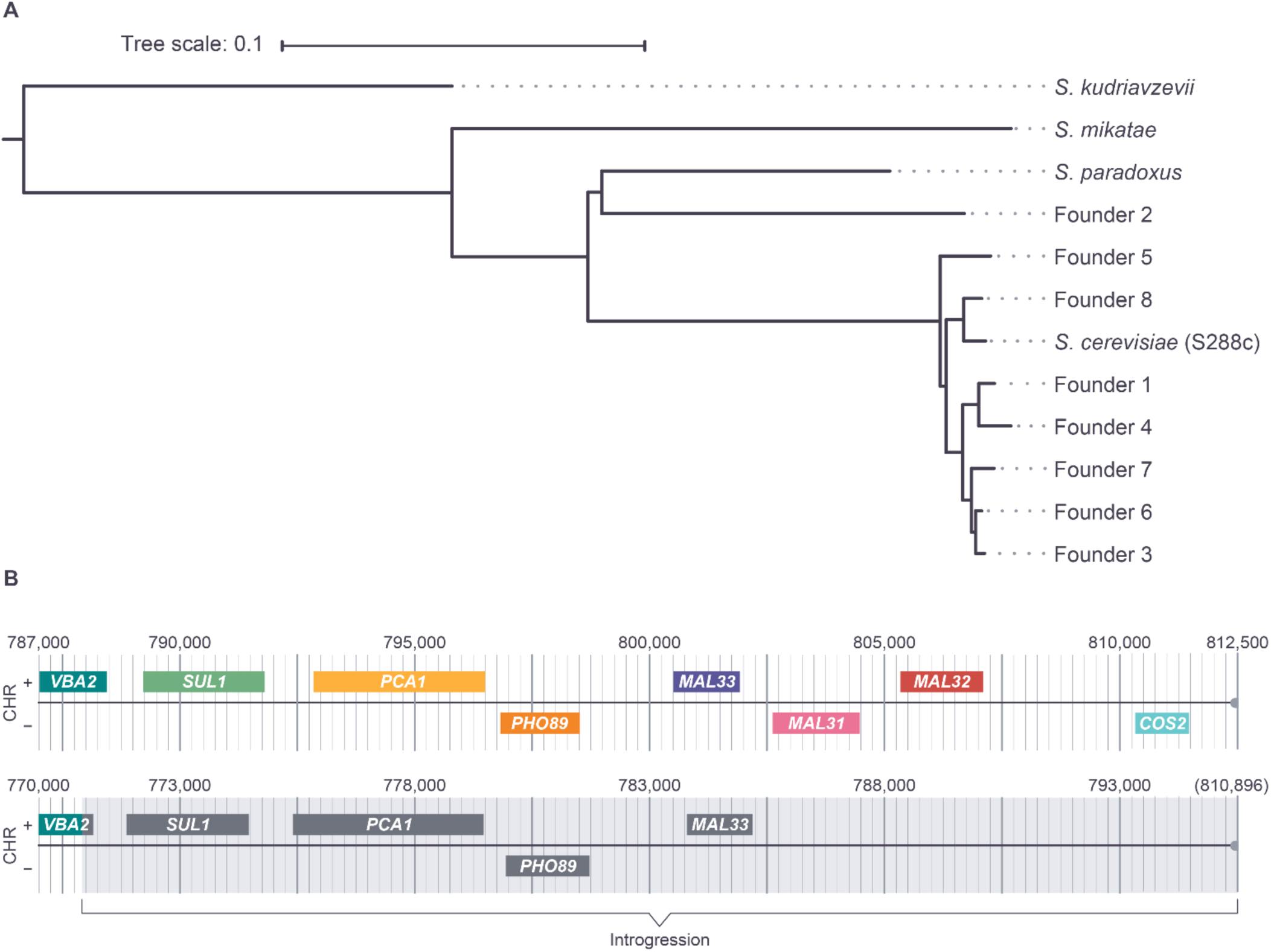
Founder 2 possesses an introgression on the right telomeric region of chromosome II that includes *PCA1*. (**A**) Neighbor joining tree of *PCA1* protein sequences. (**B**) The introgressed telomeric region of chromosome II from founder 2 (numbering from File S1) compared to the S288c reference. All ORFs >500bp displayed, with reference homolog names.

**Fig S7.**
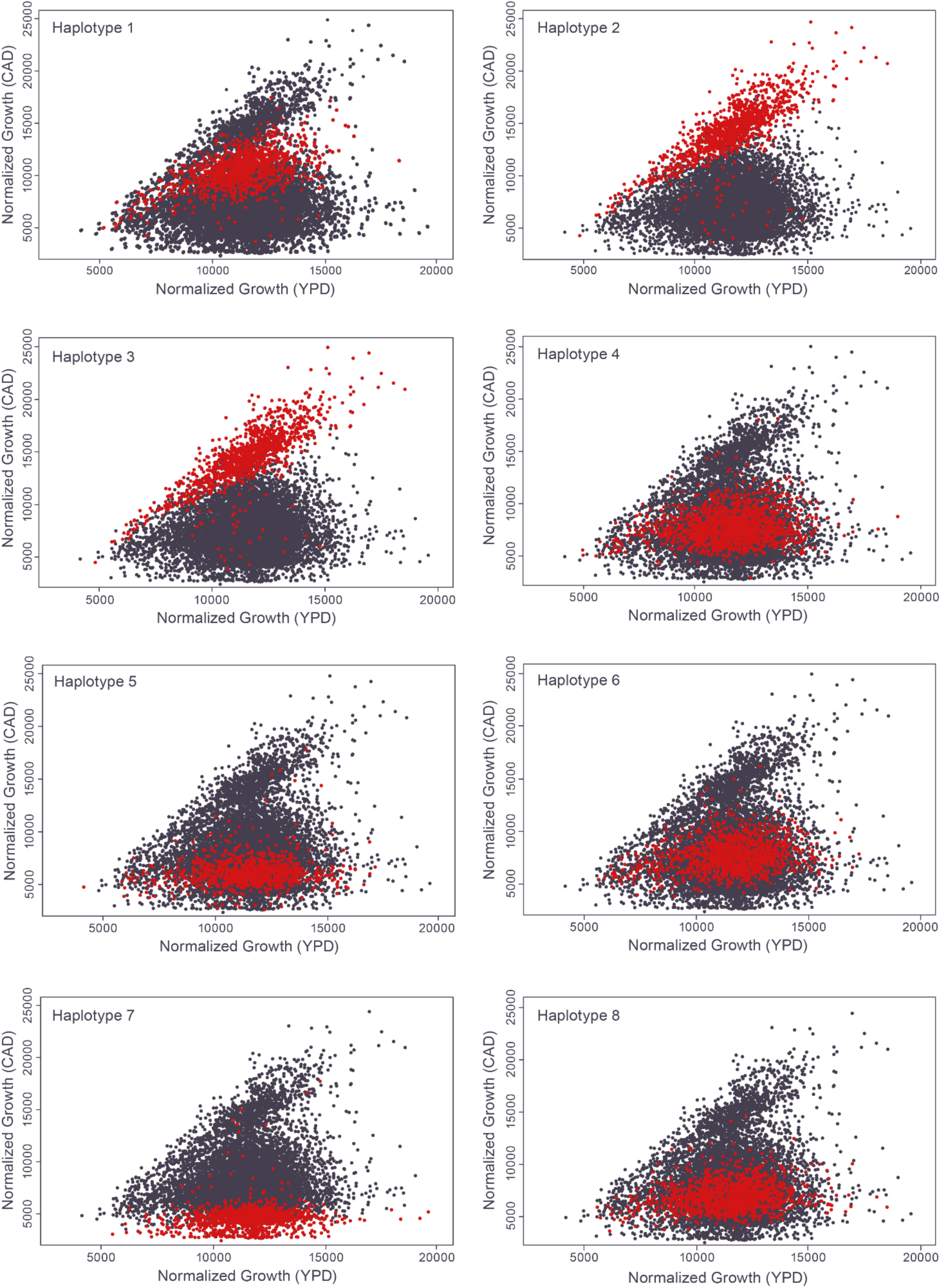
Comparisons between growth on YPD and CAD for strains stratified by *PCA1* haplotype. Full distribution of all strains shown in each plot, with current subset of strains indicated in red.

**Fig S8.**
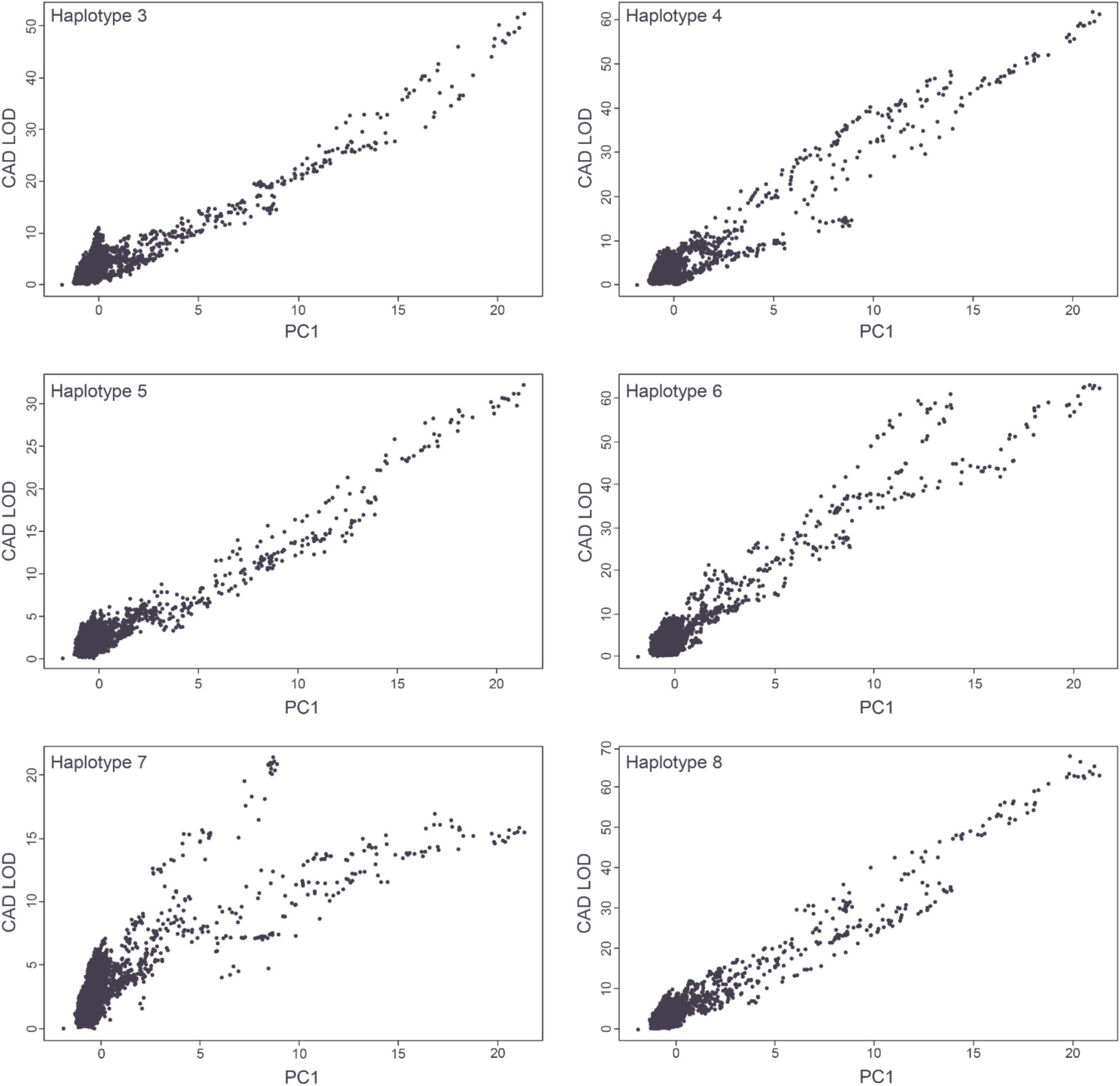
Shared Genetic Architecture on CAD for strains with *PCA1* haplotypes 3-8. LOD scores calculated at each scaffolding marker in each subpopulation plotted against first principal component of the subpopulation LODs (each scaled to unit variance).

## References

1. Timpson NJ, Greenwood CMT, Soranzo N, Lawson DJ, Richards JB. Genetic architecture: the shape of the genetic contribution to human traits and disease. Nat Rev Genet. 2018;19(2):110–24. Epub 20171211. doi: 10.1038/nrg.2017.101. PubMed PMID: 29225335.

2. Friedman JM. Implementing genomic newborn screening as an effective public health intervention: sidestepping the hype and criticism. NPJ Genom Med. 2024;9(1):70. Epub 20241219. doi: 10.1038/s41525-024-00451-7. PubMed PMID: 39702389; PubMed Central PMCID: PMCPMC11659425.

3. Xiang R, Kelemen M, Xu Y, Harris LW, Parkinson H, Inouye M, et al. Recent advances in polygenic scores: translation, equitability, methods and FAIR tools. Genome Med. 2024;16(1):33. Epub 20240219. doi: 10.1186/s13073-024-01304-9. PubMed PMID: 38373998; PubMed Central PMCID: PMCPMC10875792.

4. Bycroft C, Freeman C, Petkova D, Band G, Elliott LT, Sharp K, et al. The UK Biobank resource with deep phenotyping and genomic data. Nature. 2018;562(7726):203-9. Epub 20181010. doi: 10.1038/s41586-018-0579-z. PubMed PMID: 30305743; PubMed Central PMCID: PMCPMC6786975.

5. 5. All of Us Research Program I, Denny JC, Rutter JL, Goldstein DB, Philippakis A, Smoller JW, et al. The "All of Us" Research Program. N Engl J Med. 2019;381(7):668-76. doi: 10.1056/NEJMsr1809937. PubMed PMID: 31412182; PubMed Central PMCID: PMCPMC8291101.

6. Weiner DJ, Nadig A, Jagadeesh KA, Dey KK, Neale BM, Robinson EB, et al. Polygenic architecture of rare coding variation across 394,783 exomes. Nature. 2023;614(7948):492-9. Epub 20230208. doi: 10.1038/s41586-022-05684-z. PubMed PMID: 36755099; PubMed Central PMCID: PMCPMC10614218.

7. Valdar W, Flint J, Mott R. Simulating the collaborative cross: power of quantitative trait loci detection and mapping resolution in large sets of recombinant inbred strains of mice. Genetics. 2006;172(3):1783–97. Epub 20051215. doi: 10.1534/genetics.104.039313. PubMed PMID: 16361245; PubMed Central PMCID: PMCPMC1456308.

8. Rockman MV, Kruglyak L. Breeding designs for recombinant inbred advanced intercross lines. Genetics. 2008;179(2):1069–78. Epub 2008/05/29. doi: 10.1534/genetics.107.083873. PubMed PMID: 18505881; PubMed Central PMCID: PMCPMC2429860.

9. King EG, Merkes CM, McNeil CL, Hoofer SR, Sen S, Broman KW, et al. Genetic dissection of a model complex trait using the Drosophila Synthetic Population Resource. Genome Res. 2012;22(8):1558–66. Epub 20120410. doi: 10.1101/gr.134031.111. PubMed PMID: 22496517; PubMed Central PMCID: PMCPMC3409269.

10. Kover PX, Valdar W, Trakalo J, Scarcelli N, Ehrenreich IM, Purugganan MD, et al. A Multiparent Advanced Generation Inter-Cross to fine-map quantitative traits in Arabidopsis thaliana. PLoS Genet. 2009;5(7):e1000551. Epub 20090710. doi: 10.1371/journal.pgen.1000551. PubMed PMID: 19593375; PubMed Central PMCID: PMCPMC2700969.

11. Churchill GA, Airey DC, Allayee H, Angel JM, Attie AD, Beatty J, et al. The Collaborative Cross, a community resource for the genetic analysis of complex traits. Nat Genet. 2004;36(11):1133–7. doi: 10.1038/ng1104-1133. PubMed PMID: 15514660.

12. Lander ES, Schork NJ. Genetic dissection of complex traits. Science. 1994;265(5181):2037-48. doi: 10.1126/science.8091226. PubMed PMID: 8091226.

13. Bloom JS, Boocock J, Treusch S, Sadhu MJ, Day L, Oates-Barker H, et al. Rare variants contribute disproportionately to quantitative trait variation in yeast. Elife. 2019;8. Epub 20191024. doi: 10.7554/eLife.49212. PubMed PMID: 31647408; PubMed Central PMCID: PMCPMC6892613.

14. Boocock J, Alexander N, Alamo Tapia L, Walter-McNeill L, Patel SP, Munugala C, et al. Single-cell eQTL mapping in yeast reveals a tradeoff between growth and reproduction. Elife. 2025;13. Epub 20250312. doi: 10.7554/eLife.95566. PubMed PMID: 40073070; PubMed Central PMCID: PMCPMC11903034.

15. Parts L, Batte A, Lopes M, Yuen MW, Laver M, San Luis BJ, et al. Natural variants suppress mutations in hundreds of essential genes. Mol Syst Biol. 2021;17(5):e10138. doi: 10.15252/msb.202010138. PubMed PMID: 34042294; PubMed Central PMCID: PMCPMC8156963.

16. Otsu N, editor A threshold selection method from gray-level histograms. IEEE transactions on systems, man, and cybernetics 91; 1979.

17. Xie YJ, Q., editor A new efficient ellipse detection method. 2002 International Conference on Pattern Recognition; 2002.

18. Bates D, Mächler M, Bolker B, Walker S. Fitting Linear Mixed-Effects Models Using lme4. Journal of Statistical Software. 2015;67(1):1–48.

19. Endelman JB. Ridge Regression and Other Kernels for Genomic Selection with R Package rrBLUP. The Plant Genome. 2011;4:250–5. doi: 10.3835/plantgenome2011.08.0024.

20. Brem RB, Yvert G, Clinton R, Kruglyak L. Genetic dissection of transcriptional regulation in budding yeast. Science. 2002;296(5568):752-5. Epub 20020328. doi: 10.1126/science.1069516. PubMed PMID: 11923494.

21. Dudley AM, Janse DM, Tanay A, Shamir R, Church GM. A global view of pleiotropy and phenotypically derived gene function in yeast. Mol Syst Biol. 2005;1:2005 0001. Epub 20050329. doi: 10.1038/msb4100004. PubMed PMID: 16729036; PubMed Central PMCID: PMCPMC1681449.

22. Fisher RA. The correlation between relatives on the supposition of Mendelian inheritance. Transactions of the Royal Society of Edinburgh. 1918;52:339–433.

23. Nguyen Ba AN, Lawrence KR, Rego-Costa A, Gopalakrishnan S, Temko D, Michor F, et al. Barcoded bulk QTL mapping reveals highly polygenic and epistatic architecture of complex traits in yeast. Elife. 2022;11. Epub 20220211. doi: 10.7554/eLife.73983. PubMed PMID: 35147078; PubMed Central PMCID: PMCPMC8979589.

24. Hill WG, Goddard ME, Visscher PM. Data and theory point to mainly additive genetic variance for complex traits. PLoS Genet. 2008;4(2):e1000008. Epub 20080229. doi: 10.1371/journal.pgen.1000008. PubMed PMID: 18454194; PubMed Central PMCID: PMCPMC2265475.

25. Yang J, Benyamin B, McEvoy BP, Gordon S, Henders AK, Nyholt DR, et al. Common SNPs explain a large proportion of the heritability for human height. Nat Genet. 2010;42(7):565–9. Epub 20100620. doi: 10.1038/ng.608. PubMed PMID: 20562875; PubMed Central PMCID: PMCPMC3232052.

26. Bloom JS, Ehrenreich IM, Loo WT, Lite TL, Kruglyak L. Finding the sources of missing heritability in a yeast cross. Nature. 2013;494(7436):234-7. Epub 20130203. doi: 10.1038/nature11867. PubMed PMID: 23376951; PubMed Central PMCID: PMCPMC4001867.

27. Boocock J, Sadhu MJ, Durvasula A, Bloom JS, Kruglyak L. Ancient balancing selection maintains incompatible versions of the galactose pathway in yeast. Science. 2021;371(6527):415-9. doi: 10.1126/science.aba0542. PubMed PMID: 33479156; PubMed Central PMCID: PMCPMC8384573.

28. Adle DJ, Sinani D, Kim H, Lee J. A cadmium-transporting P1B-type ATPase in yeast Saccharomyces cerevisiae. J Biol Chem. 2007;282(2):947–55. Epub 20061114. doi: 10.1074/jbc.M609535200. PubMed PMID: 17107946; PubMed Central PMCID: PMCPMC4100611.

29. Dormer UH, Westwater J, McLaren NF, Kent NA, Mellor J, Jamieson DJ. Cadmium-inducible expression of the yeast GSH1 gene requires a functional sulfur-amino acid regulatory network. J Biol Chem. 2000;275(42):32611–6. doi: 10.1074/jbc.M004167200. PubMed PMID: 10921921.

30. McCoy RC, Wakefield J, Akey JM. Impacts of Neanderthal-Introgressed Sequences on the Landscape of Human Gene Expression. Cell. 2017;168(5):916–27 e12. doi: 10.1016/j.cell.2017.01.038. PubMed PMID: 28235201; PubMed Central PMCID: PMCPMC6219754.

31. Reilly PF, Tjahjadi A, Miller SL, Akey JM, Tucci S. The contribution of Neanderthal introgression to modern human traits. Curr Biol. 2022;32(18):R970-R83. doi: 10.1016/j.cub.2022.08.027. PubMed PMID: 36167050; PubMed Central PMCID: PMCPMC9741939.

32. Cohen JC, Boerwinkle E, Mosley TH, Jr., Hobbs HH. Sequence variations in PCSK9, low LDL, and protection against coronary heart disease. N Engl J Med. 2006;354(12):1264–72. doi: 10.1056/NEJMoa054013. PubMed PMID: 16554528.

